# Polygenic hierarchies of macroscale brain structural organisation

**DOI:** 10.64898/2026.09.19.752830

**Authors:** Yuanjun Gu, Amir Ebneabbasi, Yuankai He, Clara Pecci-Terroba, Kuldeep Kumar, Sebastian Jacquemont, Simon Baron-Cohen, HyungGyu Min, Hyejung Won, Edward Bullmore, Rafael Romero-Garcia, Richard A. I. Bethlehem, Varun Warrier

## Abstract

Different Magnetic Resonance Imaging-derived cortical, subcortical, and white matter measures are presumed to reflect different developmental and cellular processes, but how their genetic architecture is organised and the underlying cellular and developmental processes is unclear. We conducted genome-wide association studies of 2,326 imaging-derived phenotypes (IDPs) spanning twelve structural measures across the cortex, subcortex, ventricles, and white matter tracts in the UK Biobank (N_max_ = 53,751), replicated in ABCD (N_max_ = 5,119), identifying 14,176 experiment-wide significant loci and prioritizing 848 genes. We found that IDPs primarily cluster by measure into six families: brain size, cortical thickness, curvature, microstructural coherence, microstructural diffusivity, and orientation dispersion. This structure was preserved across prioritized genes, cell types, and developmental timing; brain size implicates first-trimester radial glia programs, while white matter coherence implicates adolescent astrocyte and oligodendrocyte lineages. Within individual measures, genetic effects were further organised along broad spatial gradients, with a principal gradient reflecting allometric scaling for most measures and secondary gradients showing correspondence with the sensory-association topographic axis across several measures. We identified 79 genes with spatially restricted effects along the cortex, with many aligning with these dominant gradients while a substantial subset showed additional spatial patterns. Together, these findings reveal a hierarchical organisation of the genetic architecture of macroscale human brain structure, spanning phenotype families, broad spatial gradients, and regionally restricted developmental programmes.

## Introduction

The human brain develops through a series of coordinated molecular and cellular events that shape its structure(*1*). Several of these anatomical properties can be measured using magnetic resonance imaging (MRI). These include macroscale properties such as overall size of cortical and subcortical regions, thickness and curvature of the cortex, as well as microscale properties such as neurite density and orientation. Each of these are thought to measure different underlying tissue properties. Yet, the genetic, molecular, and cellular mechanisms that give rise to these structural properties remain incompletely understood, limiting our ability to infer the molecular and cellular mechanisms underlying observable individual or group-level differences in brain structure, and their correlates with a range of neurodevelopmental, psychiatric, and neurodegenerative conditions.

A related, largely unresolved question is how genetics contribute to the organisation of the brain across and within these different measures. Genetic effects on brain structure may be principally organised by measurement modality, or by their anatomical location regardless of the measure. These two are not mutually exclusive. Distinguishing these two principles requires well-powered genome-wide association study data across multiple regions and measures.

Transcriptomic studies of the developing and postnatal human cortex have established that gene expression varies systematically across regions(*2–6*). This variation emerges from morphogen gradients in early development(*5*, *7*) and associated gradients in transcription factors alongside extrinsic inputs, notably from the thalamocortical pathways(*8*, *9*), together shaping areal identity across development. Neuroimaging studies have independently described smooth, low-dimensional spatial gradients in cortical structure and function, often dominated by the sensorimotor-to-association axis(*10*, *11*). A growing body of work has sought to connect the two, correlating imaging-derived maps against post-mortem transcriptomic atlases(*2*, *10*, *12*, *13*). Nevertheless, comparing spatial correspondence between neuroimaging and transcriptomic maps cannot fully identify causal biological mechanisms that establish the organisation, nor resolve whether distinct biological mechanisms shape the emergence and organisation of different MRI measures.

Common genetic variation offers one way to bridge this gap, by linking neuroimaging with single-nucleus transcriptomics across development. The recent availability of large-scale neuroimaging data alongside single-nucleus transcriptomic data at unprecedented scale from the developing(*14*, *15*) and postnatal human brain(*16*), provides an opportunity to integrate this with genome-wide association studies (GWAS) to identify the underlying cell types, molecular processes, and gene regulatory mechanisms that shape the brain and its topographical organisation. Recent studies have begun to conduct large-scale genome-wide association studies (GWAS) of structural properties of the cortex(*17–20*), subcortex(*21*, *22*), and white matter tracts(*23*, *24*). These studies have typically focussed on one or few cortical or subcortical measures in depth. Previous work from our team assessed the genetics of cortical structural phenotypes in up to 36,000 participants, but excluded subcortical, ventricular and tract-based measures(*20*), and was underpowered to identify significant loci for regional measures.

No study to date has combined sufficient regional statistical power with sufficient phenotypic breadth to resolve how genetic influences on brain structure are organised. We address this gap by conducting genome-wide association studies of 2,326 structural imaging derived phenotypes (IDPs) spanning cortical surface area, thickness, curvature, grey matter volume; subcortical, ventricular, brainstem, cerebellar, and corpus callosal volumes; and white matter microstructure, measured from MRI scans in up to 53,751 individuals from the UK Biobank (*25*) and up to 5,119 individuals from the Adolescent Brain Cognitive Development (ABCD) study (*26*).

## Results

### Genome-wide meta-analyses of 2,326 IDPs

We conducted Genome-Wide Association Studies (GWAS) of 2,326 cortical, subcortical, ventricular, and tract-based IDPs in up to 53,751 individuals of genetically inferred European ancestries in the UK Biobank, an increase of approximately 22,000 participants compared to a previous such comprehensive GWAS(*20*). The IDPs were derived across seven macrostructural and five microstructural measures (**Box 1**). All 12 measures were generated globally across the cortex and across 180 bilaterally-averaged cortical regions using the Glasser Parcellation scheme(*27*). Additionally, we generated GWAS of 8 subcortical volumes, two cerebellar volumes, brain stem volume, four ventricular volumes, volumes of five parcels from the corpus callosum, and five microstructural measures across 27 different white-matter tracts (**Supplementary Figure 1, Methods**).

Across all 2,326 IDP GWAS, we identified 48,540 genome-wide significant associations (P ≤ 5×10⁻⁸), of which 15,187 exceeded an experiment-wide threshold (P ≤ 4.65×10⁻¹¹, **Supplementary Table 1, Figure 1A**) derived by matrix decomposition to account for the number of independent tests conducted. SNP-based heritability was nominally significant (p < 0.05) for all IDPs (range: 0.019–0.39; **Supplementary Table 2, Figure 1B**), with limited residual population stratification based on the attenuation ratio (98.5% of all GWAS had attenuation ratio ≤ 0.2 **Supplementary Table 2**).

**Figure 1.**
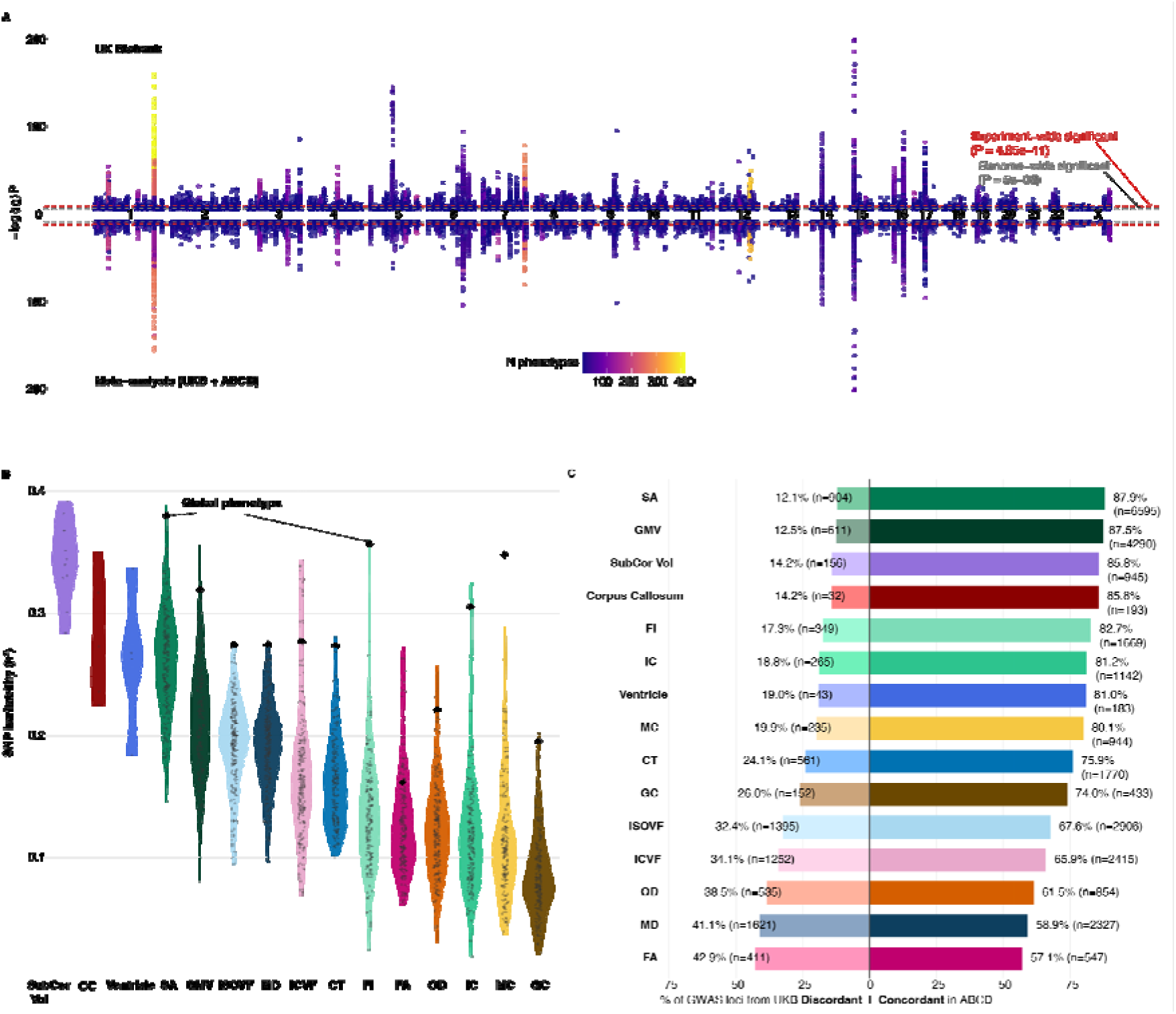
Summary of genome-wide significant findings for 2,326 imaging derived phenotypes of IDPs. **A.** Miami plot of genome-wide significant loci across all IDPs: UK Biobank discovery (top) and UKB–ABCD meta-analysis (bottom). Only lead SNPs of loci that are genome-wide significant are shown. Colour represents the number (N) of phenotypes for which a SNP reached significance. Grey dashed line, genome-wide significance (P = 5 × 10□□); red dashed line, experiment-wide significance (P = 4.65 × 10□¹¹). **B.** LDSC-based SNP heritability (h²) by phenotype. Violins, regional/tract/structure-level estimates; diamonds, global estimates. **C.** UKB–ABCD directional concordance by measure for genome-wide significant loci in the UK Biobank GWAS; bars show % concordant (right) vs discordant (left), with no of significant (n) loci labelled. Phenotypes ordered by concordance; colours as in **B**. CT, cortical thickness; SA, surface area; GMV, grey matter volume; FI, folding index; IC, intrinsic curvature; MC/GC, mean/Gaussian curvature; OD, orientation dispersion; FA, fractional anisotropy; ICVF – intracellular volume fraction, ISOVF – isotropic volume fraction; MD – mean diffusivity; SubCor Vol – subcortical volumes includes volumes of the brain stem and cerebellum; CC – Corpus Callosum

GWAS for 2,171 of 2,326 IDPs (all IDPs except tract-based) were available in 5,119 children of genetically inferred European Ancestry from the ABCD cohort (ages 9–10). Among 11,207 experiment-wide significant lead SNPs from the UK Biobank analysis that were also available in ABCD, 80.7% showed concordant effect directions and 28.32% were nominally significant (vs. null expectations of 50% and 5% respectively; **Supplementary Table 1**). Extending this to all genome-wide significant SNPs available in ABCD, (n = 35,735), 76.2% showed concordant effect directions and 19.8% were nominally significant (**Figure 1C**). Microstructural measures showed lower concordance than macrostructural measures (**Figure 1C**), consistent with continued microstructural development through adolescence and beyond(*28*). Inverse-variance weighted meta-analyses across the two cohorts (N_max_ = 58,870) identified 36,944 genome-wide and 11,529 experiment-wide significant loci. Adding loci that could not be meta-analysed with ABCD, including all tract-based IDPs, which were available only in the UK Biobank, as well as a subset of loci from other IDPs where the lead SNP was not available in ABCD, brought the total to 52,008 genome-wide and 14,176 experiment-wide significant loci, representing an approximately 3.25-fold increase compared to the previous cortex-wide systematic study (*20*) (**Supplementary Table 3, Figure 1A**). Of these, 2,639 genome-wide and 144 experiment-wide loci were not previously reported with brain imaging phenotypes. These results confirm broad replicability across different ages and cohorts. They also provide preliminary evidence for developmental sensitivity of genetic association with diverse MRI measures. We therefore focused on the GWAS generated from the UK Biobank for subsequent analyses.

#### Box 1: Measures included in the study

We use the term measure to refer to the broad phenotypic measure (e.g., surface area, volume), and imaging derived phenotype (IDP) to refer to the measure at an anatomically located parcel across global, regional, subcortical, or tract-based parcellations. We use the term family to refer to clusters of genetically correlated, parcellated IDPs from specific measures (e.g, all IDPS of GC and MC form the curvature family).

##### Macrostructural measures (T1-weighted MRI)

We quantified total cortical surface area (SA), mean cortical thickness (CT), and total cortical grey matter volume (GMV ≈ SA x CT). To characterise cortical folding, we used the two principal curvatures at each surface point (k₁ and k₂) to derive four geometric measures. Mean curvature (MC = (k₁ + k₂)/2) quantifies the average curvature of the cortex; Gaussian curvature (GC = k₁ × k₂) distinguishes dome-like (positive) from saddle-like (negative) curvature. The intrinsic curvature index (IC = max(GC, 0)) quantifies the extent of the positively curved (gyral) surface, whilst the folding index (FI = |k₁| × (|k₁| − |k₂|)) quantifies the magnitude and asymmetry of local curvature. Additionally, we measured subcortical, ventricular, and corpus callosal volumes.

##### Microstructural measures (diffusion MRI)

Diffusion-weighted imaging models water displacement in tissue. Fractional anisotropy (FA) quantifies the directional diffusion of water molecules, with 0 indicating equal diffusion across all directions, and 1 indicating diffusion confined to one axis. Mean diffusivity (MD) reflects the overall magnitude of diffusion, sensitive to membrane density and myelination. Additionally, we applied Neurite Orientation Dispersion and Density Imaging, which models water diffusion along three compartments: intracellular, extracellular and isotropic. From this model we derived intracellular volume fraction (ICVF; volume fraction occupied by neurites), orientation dispersion index (OD; degree of neurite dispersion, from aligned to isotropic), and isotropic volume fraction (ISOVF; fraction of free-water).

### MRI measure is the dominant axis of genetic organisation

To characterise the shared polygenic architecture across all 2,326 IDPs, we computed pairwise genetic correlations among all GWAS using bivariate Linkage Disequilibrium Score Regression(*29*) (**Supplementary Data 1**). We found high concordance between genetic and phenotypic correlations (r = 0.84, p < 2.2×10□¹□, Mantel test) consistent with previous findings(*20*) and Cheverud’s conjecture.

Hierarchical clustering of the genetic correlation matrix, supported by Uniform Manifold Approximation and Projection(*30*) and Leiden community detection(*31*), consistently identified six families of genetically correlated IDPs, regardless of method (**Figure 2A-D**): (i) brain size (SA, GMV, FI, IC, and subcortical volumes); (ii) cortical thickness (CT); (iii) curvature (GC and MC); (iv) microstructural coherence (FA and ICVF); (v) microstructural diffusivity (MD and ISOVF); and (vi) orientation dispersion (ODI). Clustering was driven by the type of MRI measure rather than anatomical origin of the IDP, i.e., cortical, subcortical, and tract-based IDPs derived from the same MRI measure clustered together. Although ventricular volumes and corpus callosum were distinct branches in the hierarchical clustering dendrogram, the Leiden clustering algorithm did not identify them as separate clusters, possibly reflecting their small number of phenotypes (n=4 and n=5, respectively).

**Figure 2.**
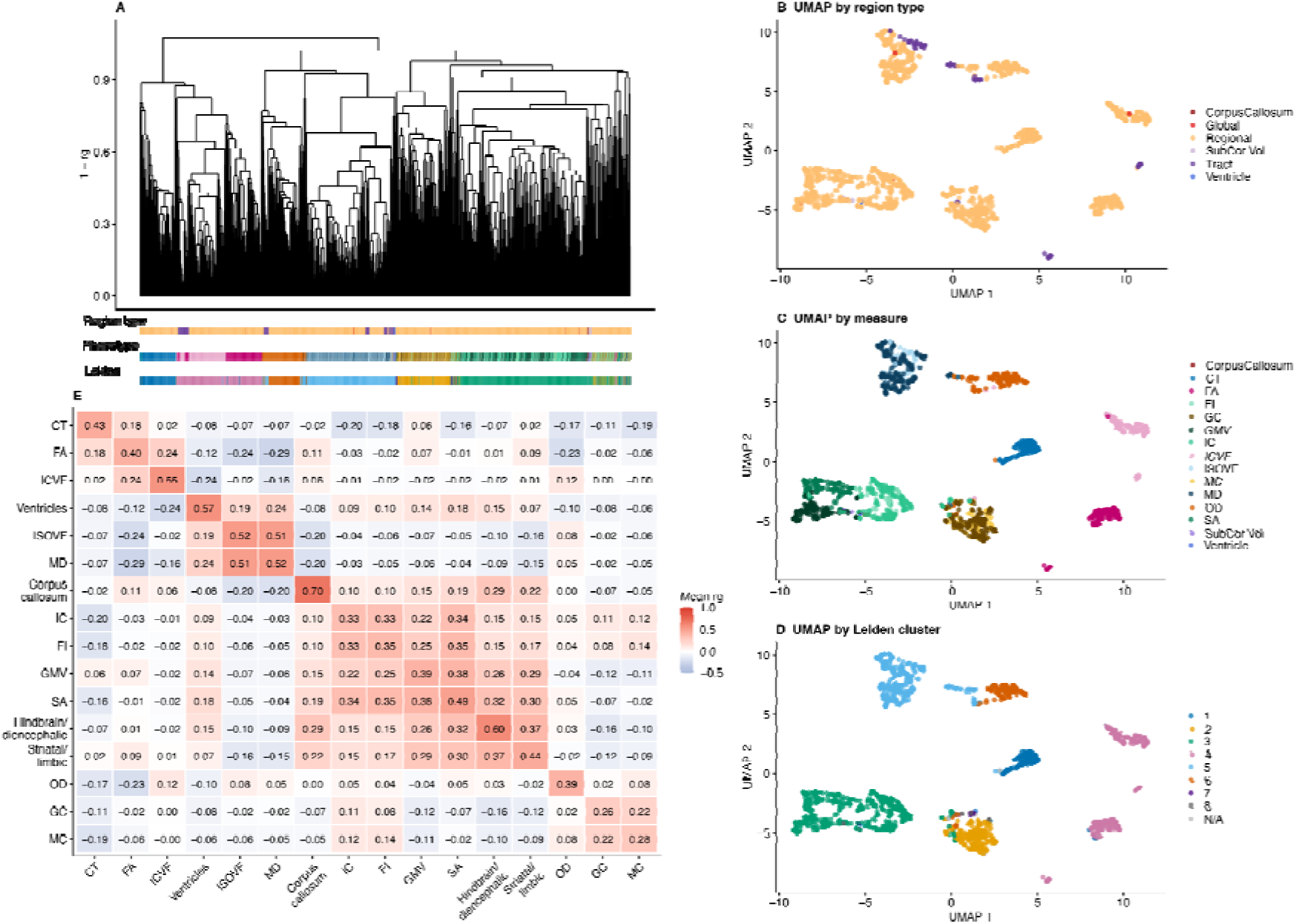
Genetic clustering of the imaging derived phenotypes. **A.** Average-linkage based hierarchical clustering dendrogram of all IDPs by genetic correlation distance (1 − r_g_), with annotation bars indicating region type (cortical, subcortical (including brain stem and cerebellum), tract, with corpus callosum and ventricles separately highlighted), IDP measure, and Leiden community membership. **(B–D)** UMAP embeddings of the same IDPs coloured by region type **(B)**, measure **(C),** and Leiden cluster membership **(D)**, demonstrating that clustering is driven by measure rather than anatomical origin. Six families are identified across methods. **E.** Mean pairwise genetic correlations between measures and IPD groups, including cortical measures and volumes of subcortical regions, ventricles and corpus callosum. Values shown are mean r_g_ across all pairwise combinations within or between groups.

Nevertheless, mean pairwise genetic correlations between ventricular volumes and corpus callosum with other measures indicated that they were outside the six-family structure (**Figure 2E**). This clustering was consistent when removing highly genetically correlated IDPs (r_g_ > 0.9 to r_g_ > 0.6, **Supplementary Figure 2**), and was further supported by LD-based clumping of the significant loci and co-localisation using Hyprcoloc (*32*) (**Supplementary Figure 3**), demonstrating that clustering was preserved both globally across the genome and when restricting to the significant loci.

Visual inspection of genetic correlations within each measure revealed additional fine-scale structure, with average within-measure genetic correlations ranging from high (ICVF, MD) to broadly dispersed (MC) (**Supplementary Figure 4A**). This fine-scale, within-measure structure is characterised in detail below.

Ventricular volumes, corpus callosum, and subcortical grey matter each showed distinct polygenic profiles (**Extended Data Figure 4B**). Ventricular volumes formed a coherent cluster. Within the corpus callosum, genetic correlations were strongest between adjacent subdivisions and weaker across the full anterior-to-posterior extent. Subcortical volumes separated into two anatomically-informed subgroups: a striatal/limbic group (caudate, accumbens, putamen, amygdala, hippocampus) and a hindbrain/diencephalic group (thalamus, pallidum, ventral diencephalon) that was genetically correlated with brainstem and cerebellum. Both these groups showed moderate positive genetic correlations with cortical brain size phenotypes.

Genetic correlations among the 27 tracts were moderately to highly positive for all five microstructural measures. The middle cerebellar peduncle and the medial lemniscus had lower genetic correlations with the rest of the tracts across all measures, likely reflecting the absence of direct cortical connectivity for these two tracts (**Extended Data Figure 4C**).

### Prioritising associated genes and biological processes

The increased statistical power of the GWAS enabled us to prioritise potential underlying genes to subsequently interrogate the underlying biology of the phenotypes. We used FLAMES(*33*) on the UK Biobank-only GWAS to prioritise genes. FLAMES prioritised 848 unique genes (10,449 gene-phenotype combinations) across all significant loci (**Supplementary Table 4, Figure 3A**). This ranged from 296 genes for SA to 71 for Gaussian curvature.

**Figure 3:**
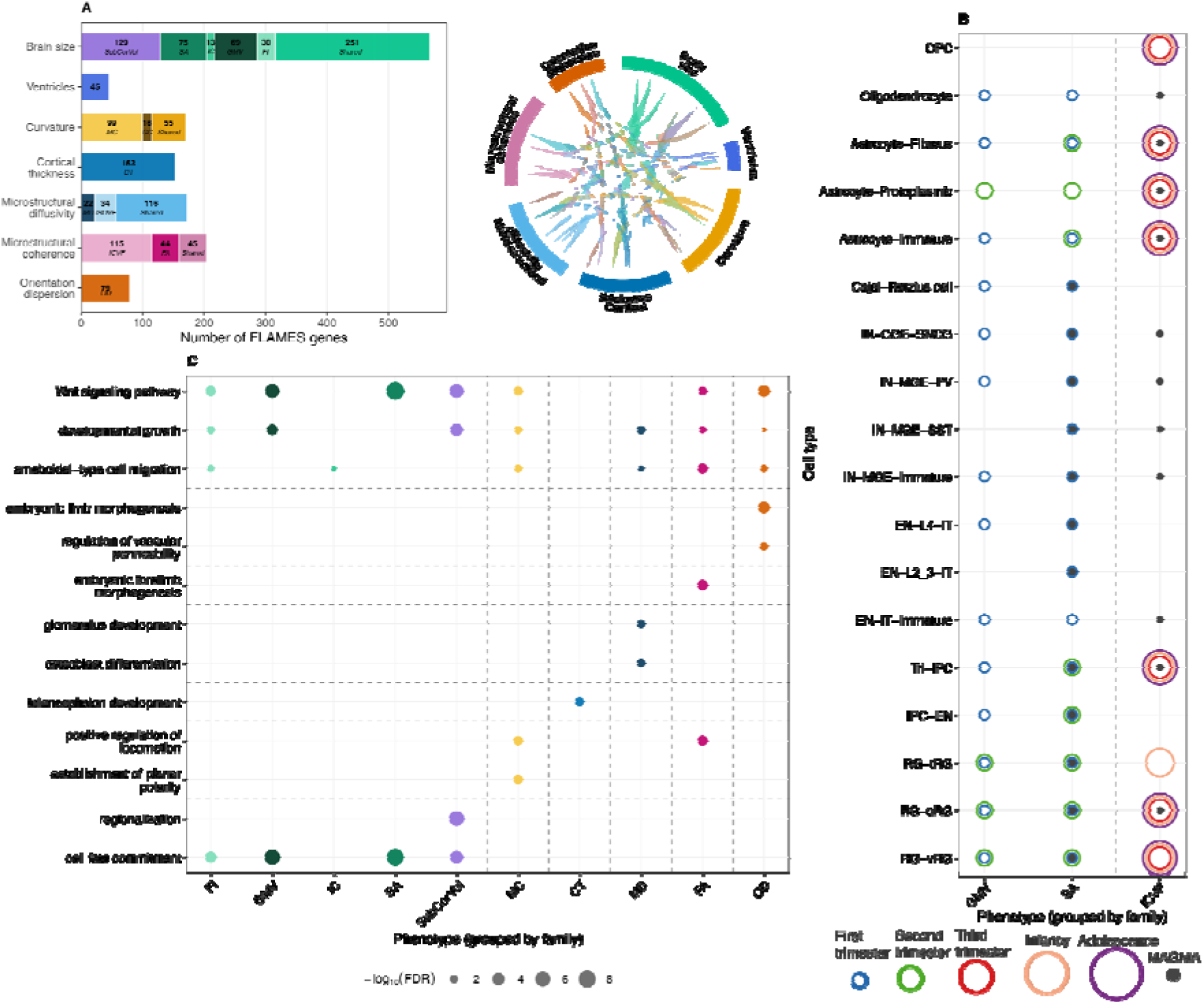
Genes identified, and cell type and gene-ontology enrichment analyses. **A.** Horizontal stacked bar chart showing the number of FLAMES-prioritised genes per phenotype family, coloured by phenotype measure (unique genes) or family colour (genes shared across ≥2 measures within a family). Chord diagram shows between-family gene overlap. **B.** Cell type enrichment of FLAMES genes across Wang 2025 fetal neocortex cell types and developmental stages. Each position shows one hollow circle per significant timepoint (logistic regression FDR < 0.05, β > 0); circle size and colour encode developmental stages. Filled dark grey inner circles indicate FDR-significant enrichment in the Wang 2025 MAGMA analysis of global GWAS signals. Phenotypes are grouped by family (dashed vertical lines). **C.** Selected Gene Ontology biological process enrichment of FLAMES genes. Each dot represents a significant term (FDR < 0.05) for a given phenotype; dot size encodes −log₁₀(FDR); dot colour encodes phenotype measure. Terms are grouped into shared (significant in ≥4 families; top, above dashed line) and family-specific (bottom, separated by dashed lines).

We used two approaches to assess the convergent validity of the genes prioritised by FLAMES. First, we leveraged regulatory variants that have been empirically validated using massively parallel reporter assays in neural progenitor cells(*34*). We detected expression-modulating variants (emVars) in close LD (r^2^ > 0.8) with 53 experiment-wide significant loci. Of these loci, 48 also had FLAMES prioritised genes. In these loci, we compared the FLAMES-prioritised genes with the genes linked to emVars using activity-by-contact scores derived from neural progenitor cells. FLAMES and activity-by-contact prioritised the same gene significantly more often than expected by chance (14 observed vs. 7.5 expected; permutation p = 0.001). Second, the FLAMES prioritized genes were enriched among genes included in clinical gene panels for cephalic disorders and malformations of cortical development (OR = 2.04, p = 9.23×10^−4^, hypergeometric test), particularly for genes linked to the brain size family (OR = 2.69, p = 6.73×10^−4^, hypergeometric test).

Consistent with findings from the genetic correlation, we observed greater overlap in the FLAMES genes identified within family than between family (Mean Jaccard index = 0.29 vs 0.15, permutation p = 1×10^−4^) (**Supplementary Figure 5**). We also found greater overlap in cell type enrichment and Gene Ontology terms within families compared to between families, demonstrating distinct biology across families (**Supplementary Figure 5**).

We investigated cell-type specificity of the families and their associated measures across prenatal and postnatal development, using cortical single-nucleus RNA sequencing (snRNA-seq) spanning the first trimester through adolescence(*14*) (**Supplementary Table 5, Figure 3B**). We used two complementary approaches: enrichment of FLAMES genes identified for all IDPs, and MAGMA(*35*)-based enrichment applied to global cortical GWAS. Brain size genes were consistently enriched in radial glia and intermediate progenitor cells across the first through third trimester, consistent with the central role of cortical neurogenesis in determining cortical expansion(*17*, *20*). ICVF and, to a lesser extent, FA (microstructural coherence family) were enriched in astrocyte populations from the third trimester through adolescence, and in oligodendrocyte populations, consistent with the protracted postnatal development of white matter microstructure and the glial origins of myelination. This enrichment for microstructural coherence in astrocytes and oligodendrocyte lineage persisted in an snRNA-seq dataset from the adult brain(*16*) (**Supplementary Figure 6**). We did not identify any significant enrichment for other families, possibly because they emerge from cell-cell interactions across multiple cell types.

We reasoned that there may be additional differences between families at the level of biological processes and cellular programmes. To identify biological processes underlying different MRI phenotypes, we conducted Gene Ontology biological process enrichment analysis of FLAMES-prioritised genes. Enrichment analyses revealed several shared biological processes: Wnt signalling, cell migration, tissue morphogenesis, and developmental growth terms were enriched across most phenotype families (**Supplementary Table 6, Figure 3C**). Additionally, macrostructural phenotypes (cortical size and curvature) shared enrichment for embryonic morphogenesis and organ development terms, consistent with progenitor-driven cortical expansion and patterning, while microstructural phenotypes (FA, ICVF, MD, OD) shared enrichment for embryonic limb morphogenesis and blood vessel morphogenesis, likely reflecting pleiotropy with mesenchymal and vascular genetic architecture. Three families showed additional distinct signals: cortical curvature was enriched for planar cell polarity terms, consistent with Wnt-planar cell polarity signalling in tangential expansion leading to gyrification(*36*, *37*); white matter diffusivity (MD) showed specific enrichment for ossification and renal system development, likely reflecting pleiotropic extra-cellular matrix architecture; and OD was specifically enriched for axon extension and positive regulation of axonogenesis, indicating that axonal cytoskeletal organisation likely contributes to OD.

### Gene regulatory mechanisms underlying brain structure

The above findings demonstrate that the families differ in their underlying genes, cell types and biological processes. These may partly arise due to differential gene regulatory mechanisms across the families. To identify this, we tested whether transcription factors (TFs) and their computationally predicted targets (eRegulons) in the developing brain(*14*) were enriched among FLAMES-prioritised genes for each phenotype (**Supplementary Figure 7A**). We identified significant enrichment for both TFs and target genes across all phenotypes, except ICVF, which may be because the dataset has poor coverage in late childhood and adulthood (**Supplementary Table 7**).

To understand the dynamic expression patterns of TFs that are also FLAMES-priorities genes, we projected these TFs along chronological time and pseudotime along the progenitor-excitatory neuron lineage. This revealed temporal differences across the families (**Supplementary Figure 7B–C**). Along chronological time, brain size and curvature related TFs (e.g, HMGA2, EMX2, PAX6, MEIS2, SOX5) peaked during the first trimester, whilst orientation diffusion and microstructural coherence related TFs peaked during adolescence, consistent with the role of astrocyte and oligodendrocytes in protracted myelination. Along the progenitor-excitatory neuron lineage, the expression of TFs associated with all families, except ODI, peaked in the progenitor stage and declined in excitatory neurons. In contrast, the mean expression of FLAMES-prioritised genes did not noticeably differ across chronological or pseudotime for any of the families. These patterns were evident when quantifying the cell-type activity (measured using AUCell) of the significant eRegulons per family. Across most families, activating eRegulons had higher mean AUCell in progenitor and macroglial cells whereas repressing eRegulons had higher activity in excitatory neurons and the oligodendrocyte lineage (**Supplementary Figure 7D**).

### Genetic topography within measures

The above analyses indicate that the primary distinction in polygenic effects in brain structure is at the level of the six families. Nevertheless, within each measure in the cortex, there is variation in genetic correlation within each measure, with mean genetic correlation ranging between 0.26 – 0.55 (**Figure 2E, Supplementary Figure 4A**). To better characterise this variation in genetic correlation, we first conducted principal component analyses of each measure’s within-measure genetic correlation matrix, and then developed a spatial-clustering framework to identify individual genes with spatially restricted effects – arealization genes (**Methods**).

Principal component analysis of the within-measure genetic correlation matrix demonstrated that the first principal component (gPC1) was dominant, explaining between 27.5% (SA) and 65.1% (CT) of variance in genetic correlation across parcels. gPC2 explained between 7.2% (CT) and 17.8% (ISOVF), and gPC3 explained between 3.3% (CT) and 16.0% (FA) of variance in genetic correlation across parcels (**Figure 4A, Supplementary Table 8**). Each measure had a differing topographic organisation across gPCs 1 – 3, with this difference becoming more distinct for gPCs 2 and 3 (**Figure 4B, Supplementary Figures 8**). Despite this, for gPCs 1 and 2 but not 3, the spatial correspondence across measures largely aligned with the family structure evident in the between-measure genetic correlations, with the exception of FI and ICI, which aligned more closely with the curvature family than the brain size family (**Figure 4C, Supplementary Figure 9**).

**Figure 4.**
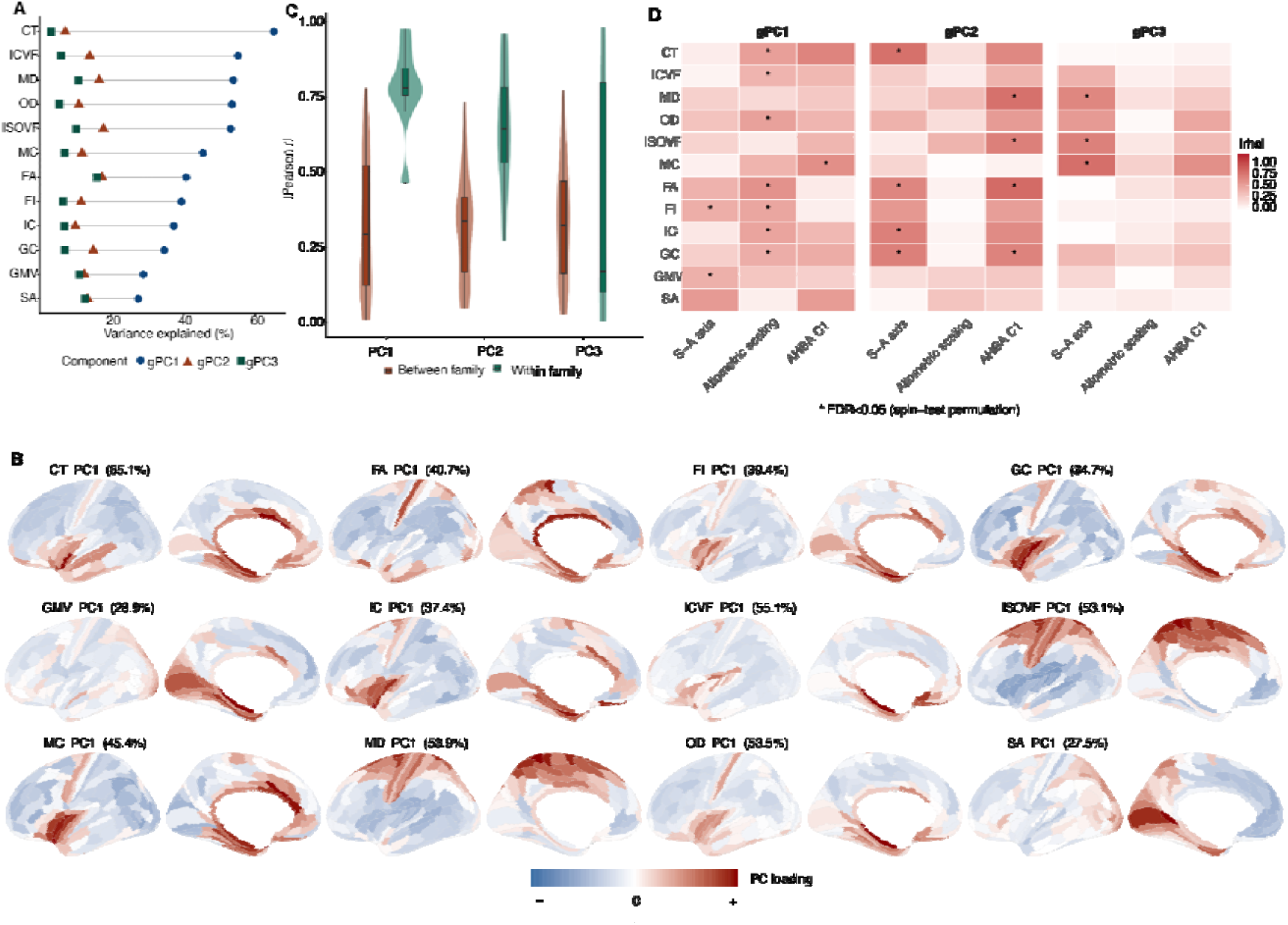
Genetic topographical organisation of the cortex. **A.** Variance explained by the first three principal components (gPC1 – 3) of the genetic correlation matrix within each cortical measure. **B.** Spatial projection of the first principal component (gPC1) on the Glasser parcellation by measure. **C.** Violin plots of within vs between family spatial correlation of gPCs 1 – 3. **D.** Enrichment of gPCs 1 – 3 in selected topographic maps.

We next tested whether each measure’s gPC1–gPC3 maps were spatially colocated with established topographical maps such as the sensory-association axis, allometric scaling and evolutionary expansion (*10*), as well as principal components of gene expression in the adult cortex (*2*), using spin-test based permutation. gPC1 colocated most consistently with allometric scaling (FDR<0.05) and the third gene expression gradient in the adult brain, but not consistently associated with the sensory-association axis. This is consistent with gPC1 predominantly reflecting a general, scaling-related axis rather than the canonical cortical hierarchy. gPC2 showed the broadest colocation with established maps, correlating moderately with the sensory-association axis, T1w/T2w myelin content, and the first gene expression principal component from the adult brain, most consistently in cortical thickness, curvature, and folding measures. gPC3 correlated most often with the sensory-association axis, particularly in diffusion and microstructural measures. Correlations were moderate in magnitude (|rho| typically 0.4 – 0.7, where significant) and measure-specific rather than universal, indicating that none of gPC1–gPC3 simply captures established topographical axes of the human cortex (**Figure 4D, Supplementary Table 9**).

### Candidate arealization genes

The gPC analysis revealed broad spatial gradients in the genetic organisation of individual measures. We next asked whether individual genes could also show more spatially restricted effects. We reasoned that genes involved in cortical arealization would be expected to show non-uniform patterns of genetic association across parcels, with stronger effects in spatially contiguous regions corresponding to distinct cortical territories. We therefore developed a spatial clustering framework to identify, from the 848 FLAMES genes, genes with spatially restricted effects, identifying 79 such genes (**Supplementary Table 10, Method**s). We refer to these as arealization candidate genes, defined by the spatial pattern of their genetic associations with regional brain structure rather than by directly measured differences in gene expression across the cortex.

Several of these arealization genes are previously reported arealization genes including: *FOXP1* (*3*, *4*), *BCL11B* (*4*), and *THBS1* (*38*) (**Figure 5A**). We also identified several novel genes which are targets of known arealization and patterning genes. This includes *MEIS1*, a transcriptional target of retinoic acid signalling which is involved in the specification of the prefrontal cortex, and whose paralog *MEIS2* is a key hub in retinoic acid gene-regulatory network in the human developing cortex (*39*). Several candidate arealization genes were prioritised for different measures. For example, *ZIC1* was an arealization gene for Volume, SA, Folding Index, and Intrinsic Curvature. For each such arealization candidate gene prioritised for two or more measures, we compared the overlap in the associated cortical parcels using Jaccard index. These parcels were more likely to overlap for pairs of measures from the same family than from different families, indicating that arealization effects also differ between families (**Figure 5B**).

**Figure 5.**
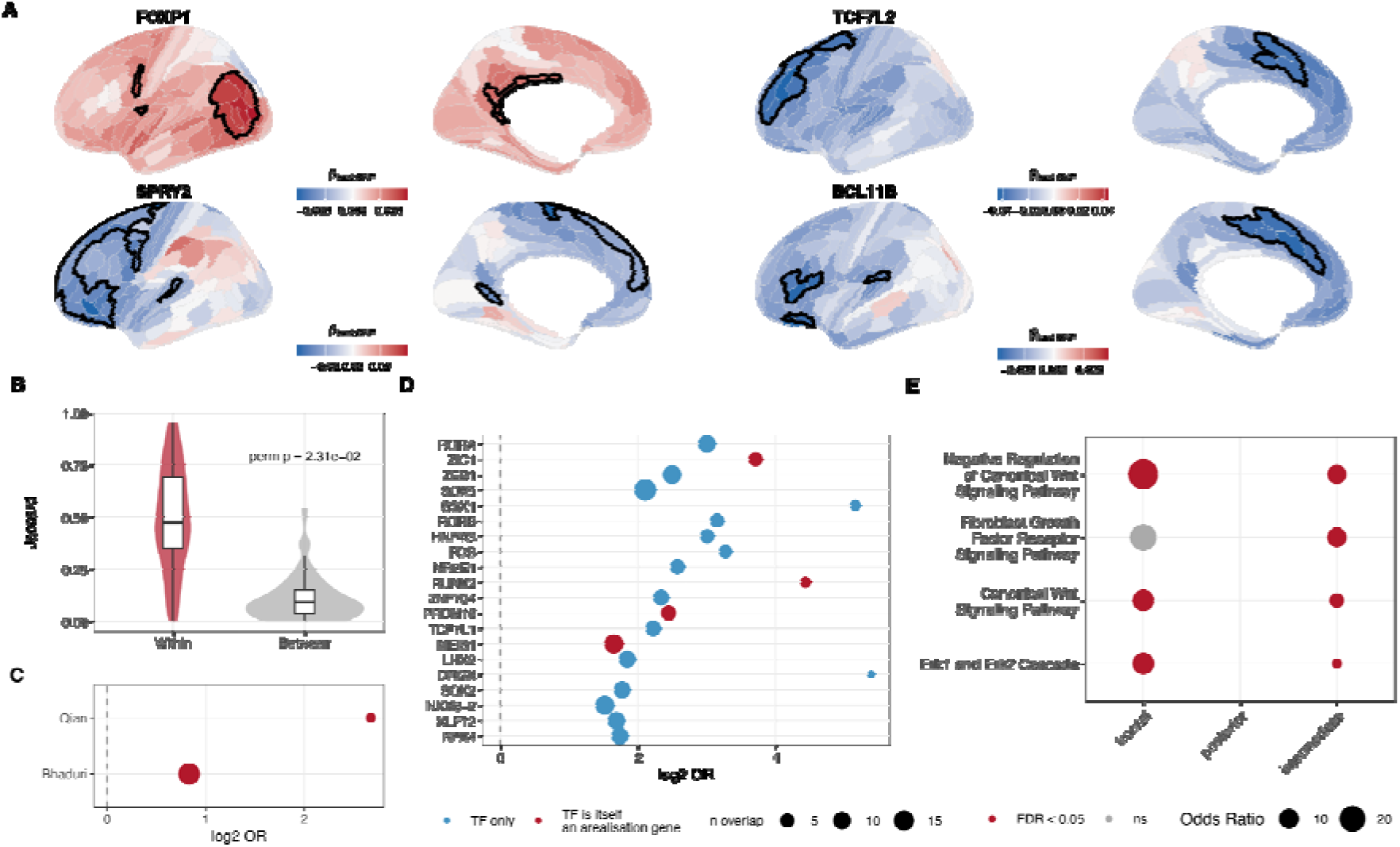
Candidate genes for cortical arealization. **A.** Cortical surface maps for four arealization candidate genes. Colour represents magnitude of lead SNP-effect in the associated loci. Outlines represent significant parcels (SA – surface area, FI – folding index). **B.** Jaccard similarity of significant parcel sets for candidate genes within and between families Violin/boxplots (median, IQR, 1.5×IQR whiskers); p-values represent permutation test. **C.** Enrichment of arealization candidates in two arealization gene sets from transcriptomic data, both of which were significant at Benjamini-Hochberg FDR < 0.05. Size indicates the number of overlapping genes. D. Enrichment of arealization genes in eRegulons. All enrichment significant, only selected enrichments provided. TFs that are known arealization genes are provided in red. **E.** Enrichment of frontal/posterior/intermediate-classified arealization genes with known telencephalic patterning gradients; one-sided Fisher’s exact test.

To determine whether these spatially-localized effects are primarily reflecting each measures top three genetic gradients (gPC1 – gPC3) or reflect additional, distinct arealization effects, we correlated the effect of each arealization candidate gene’s lead SNP across the 180 cortical parcels with gPC1 to gPC3 of the corresponding measure (spin-test permutation, **Methods**). Of the 122 (out of 123) gene-measure pairs tested, 72 were significantly correlated with gPC1 (FDR<0.05), and 14 were additionally significantly correlated with gPC2 or gPC3, with 36 gene-measure pairs not correlated with any of the first three gPCs. This indicates that, although many arealization candidate genes align with the dominant topographic genetic gradients, a substantial subset show additional spatial patterns not captured by the top three gradients of their corresponding measures (**Supplementary Table 10**).

To assess whether the candidate arealization genes are supported by external data, we tested for enrichment of these 79 genes with arealization genes identified from snRNA and spatial transcriptomic data of the developing human cortex (**Methods**). As a group, arealization candidate genes were enriched for previously identified arealization genes identified from snRNA-seq (*4*) and spatial transcriptomic data (*3*, *4*) from the developing brain (**Figure 5C, Supplementary Table 11**). Although previous studies had identified increased signatures of arealization in excitatory neurons (*3*) and to a lesser extent in radial glia(*4*), the arealization candidate genes identified in this study were not enriched for any specific developmental cell types (*14*), which may reflect biological contributions from multiple cell types or limited statistical power given the relatively small candidate gene set.

We reasoned that if arealization genes are downstream effectors of the patterning transcription factor (TFs) cascade described above, their expression should be predictable from the binding targets of known patterning TFs. We therefore tested whether arealization candidate genes are enriched for computationally predicted eRegulons (TF-target gene networks) active in the developing brain. This identified enrichment in targets of several TFs (**Figure 5D, Supplementary Table 12**) including previously implicated in arealization or patterning (e.g, LHX2 (*40*), FOXG1(*40*), EMX2(*7*), and ZIC1(*41*)). This overlap suggests that the identified arealization genes are downstream targets of TFs that are involved in arealization and patterning themselves.

The patterning TFs identified above are themselves induced by morphogen gradients along the anterior-posterior axis, as described earlier. We therefore asked whether the arealization candidate genes are also enriched for specific morphogen pathways that establish that anterior-posterior axis. We partitioned the arealization genes into three groups based on their spatial clustering (anterior, intermediate, and posterior), and tested for enrichment in gene ontology terms associated known anterior-posterior patterning pathways: canonical Wnt signalling, FGF receptor signalling and the associated ERK/MAPK cascade, and Notch signalling (*5*, *40*, *42*). The anterior– and intermediate-clustering arealization genes were enriched for (negative regulation of) canonical Wnt signalling and the ERK/MAPK cascade, while the intermediate cluster was additionally enriched for FGF receptor signalling (**Figure 5E, Supplementary Table 13**). These enrichments were absent in posterior-clustering genes. This spatial pattern is consistent with the model in which FGF interacts with and suppresses the canonical Wnt signalling from the posterior hem, thereby contributing to cortical patterning (*43*, *44*).

## Discussion

Using common genetic variants, we demonstrate that the principal genetic axis of organisation of human brain macro– and microstructure is by measure rather than anatomy (**Figure 2**). Across cortical and subcortical structures, brain structure can be resolved into six broad families. These family-based distinctions are reflected in cellular, gene ontology and gene regulatory enrichments (**Figure 3**). The brain size family are enriched for cortical progenitors across the first two trimesters and implicate key regulators of cortical progenitor identity (e.g., *HMGA2, EMX2, PAX6, MEIS2*, and *SOX5*) that are involved in progenitor pool expansion, anterior-posterior patterning, and fate specification(*45*). Consistent with prior observations, these progenitors are active in the first and second trimester of pregnancy in progenitor cells. The microstructural coherence family is associated with astrocyte and oligodendrocyte lineage programmes from the third trimester through adolescence, consistent with the protracted postnatal timeline of white matter myelination(*28*). This family was also enriched for eRegulons of TFs with known roles in oligodendrocyte specification (*SOX10, SOX8*) (*46*, *47*) and astrocyte fate commitment (*SOX9, ZBTB20*)(*48*, *49*). The role of astrocytes in microstructural coherence is underappreciated; astrocytes have wide-ranging roles in regulating myelination (*50*). Despite considerable statistical power, no other cell types were enriched for the remaining four families, indicating that their effects are likely to span multiple cell types. For example, orientation dispersion showed no single enriched cell type, yet its gene ontology signature was dominated by Wnt signalling and axon extension/axonogenesis terms. Its FLAMES-TF developmental trajectory peaked at mid-gestation, and its pseudotime profile peaked at the emergence of newborn excitatory neurons. OD was enriched in the downstream targets of MAF, a transcription factor implicated in interneuron maturation and synaptogenesis in mice(*51*), and showed higher AUCell activity in inhibitory than excitatory neurons (**Supplementary Figure 7**). We speculate that OD may therefore capture a continuum from prenatal excitatory neurite outgrowth to later inhibitory circuit maturation, potentially reflecting excitatory-inhibitory circuit assembly rather than a single cell type.

Enrichments across phenotype families were concentrated predominantly in progenitor macroglial lineages with limited enrichment for mature excitatory or inhibitory neurons. This is in contrast to most psychiatric conditions, which are primarily enriched in neuronal populations (*14*, *52–54*). These differences in cellular enrichment may partly explain the low shared genetics between structural MRI phenotypes and psychiatric conditions (*5*, *8*).

Within each cortical measure there are secondary axes of genetic organisation. The dominant principal axis aligns only moderately with allometric scaling, suggesting that this gPC1 may also capture additional sources of variation (**Figure 4**). For several of these measures, the paralimbic/medial-temporal regions constitute one end of the first principal component based on genetic similarity across all parcels. These are regions with atypical (allocortical) laminar architecture, consistent with a distinct archicortical developmental origin(*55*) of the cortex. It is possible that this atypical laminar architecture may be reflected in differential underlying genetics of this region. Furthermore, for the two curvature phenotypes alongside IC and FI, this paralimbic anchoring extends to the perisylvian and insular cortex bordering the sylvian fissure, and the regional genetics of folding in these regions may be different from those in other cortical regions. Across gPCs 2 and 3 (**Supplementary Figure 8**), several measures moderately correlate with the canonical sensory-association axis, which are influenced by both intrinsic markers as well as extrinsic cues emerging from thalamocortical connectivities(*8*). For most microstructural markers, gPC2 was also moderately correlated with the primary gradient from T1w/T2w myelin-sensitive maps(*13*). This suggests that part of the shared genetic architecture captured by gPC2 may reflect regional variation in myelination itself, rather than being specific to any single microstructural measure. The imperfect correlation may be due to the fact that the genetics primarily reflect intrinsic developmental mechanisms, or because this study is based on cross-sectional MRI measures of the adult brain, in contrast to several of these axes being developed using transcriptomic and neuroimaging data from younger individuals as structural and diffusion measures(*28*, *56*) and their regional covariance(*57*) change with age.

The measures differ in the topography of their principal axes. There is some correspondence within family especially for gPCs 1 and 2, but less so for gPC3 (**Figure 4**). Furthermore even within gPC1, IC and FI align more closely with the curvature family than the brain size family. This suggests that cortical patterning has genetic influences that are distinct from those that generate the family structure, and may reflect different developmental processes.

To further investigate these spatial effects, we developed a framework to identify genes with restricted genetic effects across the cortex, which we term arealization candidate genes (**Figure 5**). These genes were enriched for previously identified arealization genes, while also identifying additional genes not previously implicated in cortical arealization. This may reflect the greater sensitivity of population-level genetic data to subtle effects, differences between gene expression and protein abundance, or developmental and spatial effects that are not captured by existing transcriptomic datasets. The enrichment of these genes for targets of patterning transcription factors and components of anterior-posterior morphogen pathways further points to a combinatorial mechanism in which spatially distributed developmental signals establish regional identity through the coordinated action of multiple regulators.

Two genes further illustrate this point. TCF7L2 is a context-dependent effector of Wnt target gene transcription (*58*) and, in this study, a predicted target of eleven distinct arealization TFs (**Supplementary Figure 10**), consistent with the combinatorial logic wherein several patterning TFs jointly control a smaller set of downstream effectors, leading to arealization. SPRY2 is an FGF-induced negative-feedback antagonist of FGF signalling (*59*, *60*), positioning it as a component of the same anterior FGF/Wnt antagonism that is thought to pattern the anterior cortex. Both TCF7L2 and SPRY2 were themselves identified as arealization genes in this study, clustering in anterior parcels.

An important limitation is that our primary analysis is based predominantly on adult brain structure, and it therefore remains unclear how these genetic hierarchies change across development. The lower genetic correspondence observed between developmental and adult samples suggests that at least some aspects of the genetic organisation of brain structure may be developmentally dynamic. Larger longitudinal and developmental cohorts will be required to determine whether the relative contribution of global and regional genetic programmes changes across the lifespan. More broadly, our findings provide a framework for interpreting the genetic architecture of structural neuroimaging phenotypes and a resource for investigating their biological and disease relevance. We identify 52,008 genome-wide significant loci and prioritise 848 genes across 2,326 structural MRI phenotypes in up to 58,870 individuals, with summary statistics made available to facilitate further investigation of the genetic architecture of human brain structure.

Together, these findings suggest a multiscale model of the genetic organisation of human brain structure. At the highest level, genetic effects are organised according to structural and diffusion measures, reflecting different cellular and developmental programmes. Within each measure, a dominant genetic axis captures a substantial component of regional variation, with additional topographic axes aligning moderately with canonical topographic maps. At a finer spatial scale, individual genes show regionally restricted effects, many of which align with these broader gradients while others are not captured by the dominant axes. These genes with spatially restricted effects that are known arealization genes, downstream targets of morphogens or patterning TFs underlie these genetic gradients.

## Methods

### Participants

We included any individuals who have both genetic and neuroimaging data from the latest UK Biobank neuroimaging cohort (UKB) (*25*) and the Adolescent Brain Cognitive Development (ABCD) cohort(*26*). We restricted the participants to participants of self-reported European ancestries, and further excluded participants with excess homozygosity and sex mismatch, and those who were more than *±*5 SD deviated from the mean of the first two genetic principal components. We also excluded participants who deviated more than *±*5 SD/MAD from mean and median respectively of the scaled MRI-derived phenotypes, resulting in a maximum of 53,751 participants in the UKB and 5,119 participants in ABCD who were included in the GWAS.

### MRI Acquisition and Preprocessing

#### UK Biobank

For the MRI analyses, preprocessed T1– and diffusion-weighted MRI minimally processed data were obtained from the UK Biobank (application 20904). T1-weighted structural images were acquired using a 3D sagittal MPRAGE sequence at 1 mm isotropic resolution (208 × 256 × 256 matrix) with in-plane acceleration (iPAT = 2), prescan normalization, and a scan duration of five minutes. Diffusion-weighted images were acquired using a spin-echo echo-planar imaging (SE-EPI) sequence at 2 mm isotropic resolution (104 × 104 × 72 matrix). The protocol included five b = 0 images (plus 3 reverse phase-encoding b = 0 images), 50 diffusion-weighted volumes at b = 1000 s/mm², and 50 at b = 2000 s/mm², with 100 unique diffusion-encoding directions. Imaging used a monopolar Stejskal–Tanner diffusion preparation (TE = 92 ms; δ = 21.4 ms, Δ = 45.5 ms) with multiband factor 3 and a total acquisition time of seven minutes.

T1-weighted images were further processed using FreeSurfer 6.0.0 (*61*), with T2-FLAIR images incorporated when available to refine pial surface reconstruction. The FreeSurfer recon-all pipeline generated image-derived phenotypes through automated intensity normalization, skull stripping, surface reconstruction, topology correction, and cortical parcellation. Subcortical structures were segmented using FreeSurfer’s aseg tool, with additional FreeSurfer sub-segmentation performed for selected structures where available.

Diffusion MRI data were corrected for eddy currents and head motion, with slice-wise outliers corrected using FSL’s eddy, followed by gradient distortion correction(*62*). Diffusion tensor metrics, including fractional anisotropy (FA) and mean diffusivity (MD), were estimated from the b = 1000 s/mm² shell using DTIFIT. In parallel, neurite microstructure was modelled using NODDI implemented through the AMICO framework(*63*), yielding intracellular volume fraction (ICVF), isotropic volume fraction (ISOVF), and orientation dispersion index (OD). In addition, tractography-based analysis was performed by modelling within-voxel fibre orientations using bedpostx, followed by probabilistic tractography with crossing-fibre modelling using PROBTRACKX. Twenty-seven major white matter tracts were automatically reconstructed using the AutoPtx framework, and tract-specific diffusion measures were computed as tractography-weighted mean values of diffusion tensor and NODDI-derived metrics.

#### ABCD

High-resolution structural imaging included a 3D T1-weighted (T1w) inversion-prepared RF-spoiled gradient echo sequence and a 3D T2-weighted (T2w) variable flip angle fast spin echo sequence, both acquired at 1 mm isotropic resolution. Prospective motion correction was applied to the T1w and T2w acquisitions when supported by the scanner. Diffusion MRI data were acquired at 1.7 mm isotropic resolution using a multiband echo-planar imaging sequence with a slice acceleration factor of 3. The protocol included 96 diffusion-weighted directions, seven b = 0 images, and four diffusion shells (b = 500, 1000, 2000, and 3000 s/mm²), comprising 6, 15, 15, and 60 directions, respectively. Acquisition protocols were harmonized across Siemens, GE, and Philips scanners to maximize cross-site consistency.

Structural MRI preprocessing included correction for gradient nonlinearity distortions, registration of T2-weighted images to T1-weighted images, correction for intensity inhomogeneity (bias-field correction), rigid-body alignment to a standard reference space, and extensive automated and manual quality control. Cortical reconstruction and subcortical segmentation were performed using FreeSurfer 7.1.1.

Diffusion MRI data were corrected for eddy current-induced distortions, participant head motion, motion-related signal dropout (slice-wise outliers), susceptibility-induced (B0) distortions using reverse phase-encoding images, and gradient nonlinearity distortions. Diffusion images were registered to the T1-weighted structural image, and diffusion gradient directions were reoriented following motion correction. Diffusion tensor modelling, NODDI, and tract-based analyses were then performed using the same procedures as those described for the UK Biobank. For a full list of IDPs studied, please refer to **Supplementary Table 14**.

### Generation of MRI Phenotypes

The processed MRI-derived imaging data was parcellated and aligned against the Human Connectome Project parcellation (i.e. Glasser parcellation)(*27*), which generated 360 parcels spanning the two hemispheres. Parcellation was done via surface-to-surface registration against the HCP fsaverage template and in the case of the microstructure measure with the additional intermediate step of volume to surface projection (https://github.com/ucam-department-of-psychiatry/UKB). Regional cortical MRI-derived phenotypes were generated by averaging across the hemispheres, and global cortical phenotypes by averaging or summing across all regions. Across these cortical global and regional parcels, we generated 12 imaging phenotypes: 1. Surface area, 2. Grey-matter volume, 3. Folding Index, 4. Intrinsic Curvature Index, 5. Gaussian Curvature, 6. Mean Curvature, 7. Cortical thickness, 8. Fractional Anisotropy, 9. Mean Diffusivity, 10. Intracellular Volume Fraction, 11. Isotropic Volume Fraction, 12. Orientation Dispersion Index. The first seven are macrostructural phenotypes, whilst the last five are diffusion-based microstructural phenotypes. Additionally, we included volumes of four ventricles, eight subcortical regions, two cerebellar regions, brain stem, and five corpus callosal parcels.

### Genetic Data Preprocessing and Quality Control

The pipeline used for quality control and imputation (HRC + UK10K/1kGP) of the genotype data is provided in detail elsewhere for UKB(*64*). For ABCD, genotypes were imputed using the TOPMED imputation server, details of which are provided elsewhere(*65*). Post-imputation, we excluded SNPs: (1) that were multiallelic; (2) without unique rsids; (3) with imputation scores (r^2^) < 0.4; (4) with genotyping rate < 0.9; (5) with minor allele frequency < 0.1%; (6) not in Hardy-Weinberg equilibrium (P<1E-6). For sex chromosomes (X and Y), we only included SNPs from the pseudo-autosomal regions. This resulted in a list of 8,658,529 SNPs that were used for the subsequent GWAS analyses. In ABCD, additionally, we included only SNPs present in UKB given the difference in sample sizes and statistical power between the two cohorts.

### Genome-wide Association Study and meta-analyses

In the UK Biobank, we conducted GWAS for all phenotypes generated for autosomal and X chromosome using GCTA FastGWA (v1.93)(*66*). We used a mixed linear model and included age, age^2^, sex, age × sex, age^2^ × sex, genotype batch number, MRI scanning site, first 40 genetic principal components, mean framewise displacement, maximum framewise displacement, and Euler Index as covariates as previously done(*20*). In addition, we also include the availability of T2 scans as discrete covariates for all structural IDPs. Regional GWAS for FI at the ‘AVI’ region was excluded from downstream analysis, due to low residual variance.

For ABCD, we used summary statistics from previously conducted GWAS of global and regional phenotypes(*20*). Additionally, we generated GWAS for subcortical, cerebellar, brain stem, corpus callosal, and ventricular volumes using the same pipeline as that for UKB. Inverse-variance weighted meta-analyses was conducted in PLINK 1.9(*67*) to meta-analyse the GWAS summary statistics of ABCD and UKB.

For both the UKB-only and the meta-analysed GWAS, we defined genome-wide significant threshold at 5×10^−8^; and an experiment-wide significant threshold of 4.65 × 10^−11^. This was identified by matrix decomposition(*68*) of the phenotypic correlation matrix to identify 1,075 independent IDPs followed by Bonferroni correction (5E-8/1,075).

### Identification of novel loci

We identified significant loci from sixteen previous neuroimaging studies spanning structural, diffusion, and functional phenotypes(*17*, *19–24*, *69–77*). Lead SNPs reaching genome-wide significance in this study were compared against previously identified lead SNPs, and were considered novel if they were not within 50kb distance and not in LD (r² ≥ 0.1, 250kb window) against a reference panel of 5,000 UK Biobank participants.

### Clumping and co-localisation

We clumped all loci that reached the experiment-wide significant threshold (*r*^2^ = 0.1; 1,000 kb window) to identify significant loci, using LD from a subset of 5,000 unrelated individuals from the UK Biobank included in the GWAS.

We used HyPrColoc (v1.0)(*32*), a Bayesian method that enables multi-trait colocalization analysis and is robust to participant overlap, to identify co-localised loci across all GWAS in the UK Biobank, restricting to experiment-wide significant loci. We mapped these onto predefined approximately independent LD blocks derived from the 1000 Genomes European population (GRCh37/hg19) as implemented in the mapgen R package(*78*). HyPrColoc was then applied using default prior probabilities and inputs. We identified co-localised genomic regions using a posterior probability threshold of 0.95, ensuring high confidence in shared causal variants across traits. The branch-and-bound divisive clustering algorithm implemented in HyPrColoc was used to identify clusters of phenotypes sharing causal variants at each locus.

### SNP-based heritability, genetic clustering and outlier detection

SNP-based heritability for both the UKB MRI GWAS and the meta-analysed MRI GWAS, as well as the genetic correlations between all UKB MRI GWAS, were estimated using LDSC (v1.01)(*29*, *79*), with LD scores based on European-ancestry reference data, without constraining the intercepts. From the LDSC-derived genetic correlation matrix, we conducted hierarchical clustering (average linkage) of the genetic dissimilarity matrix to identify clusters. This was supported by the Leiden community detection algorithm(*31*) after rescaling the genetic correlation to 0 – 1 ((rg + 1)/2). Clusters containing fewer than 10 traits were grouped into an “unassigned” category. The results of the different clustering methods were then visualised through Uniform Manifold Approximation and Projection (UMAP)(*30*).

### FLAMES analyses

Gene prioritisation was performed using FLAMES (Fine-mapped Loci Assessment Model of Effector genes)(*33*), a machine learning framework that integrates GWAS summary statistics, fine-mapping results, and functional genomic annotations to prioritise likely causal genes. Gene-based association testing was first conducted using MAGMA (v1.10)(*35*), with SNPs assigned to genes by physical proximity using PoPS reference annotations (magma_0kb.genes.annot, generated February 16, 2021)(*80*) rather than more recent MAGMA annotations, to ensure compatibility with FLAMES pre-computed features. PoPS feature scores were generated using pathway-naïve annotations. Fine-mapping used 95% credible sets constructed via approximate Bayes factor (ABF) methods (coloc v5.2.3)(*81*), which does not assume a single causal variant per locus. FLAMES was then applied and a 75% cumulative precision threshold used to define the high-confidence prioritised gene set for each phenotype, corresponding to genes meeting both a scaled FLAMES score > 0.248 and a raw FLAMES score > 0.136. Phenotypes were excluded where no SNPs reached experiment-wide significance thresholds for fine-mapping, or where no prioritised genes met both threshold criteria.

#### Validation of FLAMES genes

##### Analyses with expression-modulating variants

We reanalyzed previously reported MPRA data for variants associated with cortical structure GWAS traits(*34*) to identify expression-modulating variants (emVars), following our established analysis framework(*82*). First, allele-specific activity was assessed by comparing the effect allele (A1) with the non-effect allele (A2) using mpralm(*83*), with replicate effects included in the model. P-values were adjusted for multiple testing using the Benjamini-Hochberg procedure, and variants with a false discovery rate (FDR) < 0.1 were considered to show significant allelic differences. Second, enhancer activity was assessed by comparing each allelic construct with a scrambled-sequence control using a mixed-effects model implemented in lme4 (https://www.jstatsoft.org/article/view/v067i01), with replicate included as a random effect. For each variant, the allele with the smaller P-value was used for multiple-testing correction, as previously described(*82*). Variants with an FDR < 0.1 and log2 fold change (log2FC) > 0 were considered to exhibit enhancer activity. Variants meeting both the allele-specific activity and enhancer-activity criteria were classified as emVars(*82*). emVars were matched to GWAS lead SNPs by direct position overlap or linkage disequilibrium (r² > 0.8, European reference panel, LDlinkR)(*84*), within a 500-kb window of each lead SNP. This resulted in 53 experiment-wide significant loci wheret the lead SNP overlapped or was in linkage disequilibrium with an emVar.

To identify putative target genes of emVars, we first generated candidate emVar-gene pairs by selecting all protein-coding genes whose promoters were located within ±500 kb of each emVar. For each candidate pair, we quantified Micro-C–based chromatin contact frequency in neural progenitor cells (NPCs) between the 10kb bin containing the emVar and the 10kb bin overlapping the gene promoter. When a promoter overlapped multiple bins, the maximum contact count across those bins was used. Chromatin accessibility at each emVar was quantified as the mean NPC ATAC-seq signal within a 500bp window centered on the variant(*85*). Using the chromatin contact frequency and ATAC-seq signal, we calculated an activity-by-contact (ABC) score for each candidate emVar-gene pair as: log(Micro-C contact + 1) x log(ATAC signal + 1). ABC scores were then standardized as Z-scores across all candidate emVar-gene pairs within each locus. Candidate pairs with a locus-specific ABC Z > 1.5 were retained as the final emVar-gene mappings, following our previously reported approach(*82*).

Of the 53 experiment-wide lead SNPs which were in high LD with an emVar, 48 had both a FLAMES prioritised gene as well as an MPRA-ABC-prioritised gene. For each of these lead variants, we defined a 500-kb window, merging overlapping windows. We then identified all protein-coding genes overlapping this window. For each of the 48 lead SNPs, a gene was concordant when it was prioritised by both MPRA-ABC and FLAMES. Enrichment for concordant genes was assessed using within locus permutation testing (10,000 permutations). In the permutation tests, the number of MPRA-ABC-prioritised genes at each locus was held fixed, and gene identity was randomly resampled without replacement from that locus’s candidate gene pool to create a null distribution, against which the observed number of concordant genes were compared.

We additionally also investigated overlap with genes associated with cortical malformations (e.g., severe microcephaly, lissencephaly) using genes from the “Malformations of Cortical Development (v8)” and “Severe Microcephaly (v8)” panels from Genomics England(*86*) and the ClinGen(*87*) Expert Panel on “Brain Malformations”. For the latter, we restricted it to only genes with the “Definitive” classification. Enrichment was done using a hypergeometric test, with all protein coding genes as background.

### Cell type enrichment

Cell type enrichment of FLAMES-prioritised genes stratified by the twelve cortical measures was assessed using single-nucleus RNA-sequencing data from two reference datasets: the Wang 2025 fetal neocortex atlas that spanned the first trimester to adolescence(*14*), and the Siletti 2023 adult human brain atlas(*16*).

For each reference dataset, a pseudobulk expression matrix was constructed by computing mean log-normalised expression per cell type (and developmental stage for Wang). The background gene set comprised all protein-coding genes detected in the respective pseudobulk. For each measure and each cell type × developmental stage bin (Wang) or cell type (Siletti), enrichment was assessed via logistic regression:

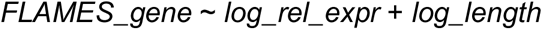

where *log_rel_expr* is the log□□-transformed expression in the target bin minus the gene’s mean log□□ expression across all bins. We corrected for multiple testing using Benjamini–Hochberg FDR correction.

Additionally, we conducted MAGMA based enrichment of developmental(*14*) and adult cells(*16*) for the twelve global cortical GWAS using cell-type specificity scores derived from Cepo(*88*). Significant enrichment was identified by applying Benjamini-Hochberg FDR correction within each measure across all cell types.

### GO biological process enrichment

Gene Ontology (GO) biological process enrichment of FLAMES-prioritised genes was conducted per phenotype using over-representation analysis implemented in clusterProfiler (*89*)(v4; enrichGO, org.Hs.eg.db). The background gene set comprised all protein-coding genes in the Wang 2025 pseudobulk. Significant terms (Benjamini-Hochberg FDR□<□0.05) were simplified using semantic redundancy reduction (clusterProfiler::simplify, similarity cutoff□=□0.7). Additionally, for visualisation, GO terms with pairwise similarity ≥□0.5 as per the Wang method(*90*) implemented in GOSemSim(*91*) were removed, retaining the term with higher mean −log₁₀(FDR) across families.

### Family similarity analyses

We conducted a series of analyses to quantify similarity between measures within and across families. FLAMES gene overlap between measures was quantified using the Jaccard index. Cell type enrichment profile similarity was quantified as the Pearson correlation between measure-level β-coefficient vectors across the cell-type x developmental stage bins. GO enrichment profile similarity was quantified as the Spearman correlation between −log₁₀(FDR) vectors across the union of all tested GO terms, with missing terms assigned a score of zero.

For each similarity measure, the difference between mean within-family and mean between-family pairwise similarity was calculated. The significance of this difference was calculated by permuting the family assigned to each measure and recalculating the difference. This was done by shuffling family-measure labels 10,000 times. Additional leave-one-out analyses were conducted to verify whether these results were driven by the Brain Size family.

#### Enrichment for Transcription Factors and eRegulons

We next assessed whether the FLAMES genes, stratified by measure, were enriched for transcription factors and their predicted target genes (eRegulons) identified by SCENIC+(*92*) using the Wang et al., 2025 data(*14*), and p-values were adjusted for multiple testing via Benjamini-Hochberg FDR correction separately for transcription factors and target genes. The enrichment was done using logistic regression:

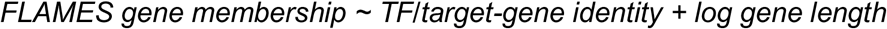

For significant TFs, we calculated the mean expression (based on the Wang 2025 data) of TFs within measure, normalised across all genes within each developmental timepoint to identify the measure-specific trajectory across chronological time and across progenitor-excitatory neuron lineage-based pseudotime. To contextualise this, we conducted the same analyses across all FLAMES genes. We then calculated the normalised (z-scored) regulon activity (AUCell) for each eRegulon across cell types, averaged within family, regulatory polarity (activating/repressing), and broad cell type, to summarise eRegulon activity patterns across the neurodevelopmental cell-type hierarchy.

### Principal component analysis of within-measure genetic correlation

We used the within-measure genetic correlation matrix and extracted the first three principal components (gPC1-3) from the associated dissimilarity matrix (1 – rg). Spatial correspondence of gPC maps within and between measures was quantified using the Pearson correlation of parcel-wise loadings, with within-family versus between-family differences tested using the same permutation framework described above (Family similarity analyses). Spatial correspondence was also tested with seven topographic maps: the first three components of AHBA transcriptomic variation(*2*), and maps of allometric scaling, evolutionary cortical expansion, the sensory-association axis, and the T1w/T2w myelin-sensitive ratio(*10*). Correspondence was assessed using 10,000 spin-test based permutation(*93*). Observed correlations were compared against this null to derive a spin-test p-value (p_spin), and p-values were corrected for multiple comparisons using Benjamini-Hochberg FDR within each measure × gPC combination.

The same spin-test procedure was used to test whether each arealization candidate gene’s lead-SNP effect (|beta|) across the 180 cortical parcels was spatially correlated with that measure’s gPC1, gPC2, or gPC3 loadings (see Identifying arealization candidate genes, below).

### Identifying arealization candidate genes

To identify arealization genes from the FLAMES prioritised genes, we first identified FLAMES genes that were significant for at least 9 cortical parcels (5% of 180 bilateral Glasser parcels), per cortical measure. We computed a spatial clustering score as the ratio of observed adjacent parcel pairs (i.e., parcels with a shared boundary) among the significant parcels to the expected number under a permutation null, by permuting the significant number of parcels per arealization candidate gene 1000 times. This preserved the number of significant parcels while randomising their cortical locations. Benjamini-Hochberg FDR correction was applied across all gene × phenotype combinations on the permuted p-values. We then used three additional filters per arealization gene. First, we restricted it to genes that were significant in only up to 60 parcels (one third of all parcels), as genes significant in more parcels may be capturing global cortical effects rather than arealization effects. Second, we required at least 30% of all significant parcels for the arealization candidate genes to be contiguous. This would remove genes that are significant across scattered parcels. Third, we counted how many separate, disconnected groups of significant parcels each gene had (where a group is one or more parcels connected to each other by a shared border), and excluded genes with more than 8 such groups. This excludes genes that have one reasonably large contiguous patch (satisfying the second filter above) but whose remaining significant parcels are scattered across many small, disconnected fragments elsewhere in the cortex. Finally, for all arealization genes that passed this quality filter, we tested whether the associated lead SNP’s effect was spatially concentrated within the gene’s own significant parcels, using two complementary tests: (i) a Wilcoxon rank-sum test comparing |beta| inside versus outside the parcels associated with the arealization candidate gene, and (ii) a spin-test-based Spearman correlation between |beta| and geodesic distance from the nearest associated parcel, testing for the expected negative (distance-decay) relationship. Three gene-measure pairs (WNT3/SA, RUNX2/MD, KANSL1/GMV) failed to show this pattern under either test and were excluded from all downstream analyses, yielding a final set of 79 arealization candidate genes across 123 gene-measure pairs.

### Enrichment analyses of arealization candidate genes

We tested whether the arealization genes were enriched for two previously identified sets of arealization genes(*3*, *4*) from transcriptomic data of the developing postmortem cortex using one-sided Fisher’s exact test, with all protein-coding genes as the background. We then tested whether arealization genes were enriched for cell-type specific expression in developmental stage bins in the Wang (2025)(*14*) dataset using this formula:

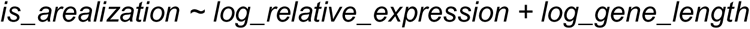

Using eRegulons identified from Wang et al., (2025) we investigated whether the arealization genes were enriched for eRegulons using this formula:

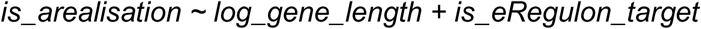

For both the cell-type x developmental stage and the eRegulon enrichment, we used the genes expressed in the Wang dataset as background.

Finally, we investigated whether arealization genes were enriched for well-characterised anterior-posterior (AP) patterning pathways, and for regulatory targets of transcription factors with established roles in cortical arealization.

We first classified each arealization gene as frontal, posterior, or intermediate based on at least 50% its significant parcels falling within a predefined frontal parcel set or a predefined posterior parcel set (both derived from the Glasser parcellation). Other genes were classified as intermediate. For each cluster, we tested for enrichment against six GO Biological Process gene sets (MSigDB C5) reflecting established anterior-posterior patterning pathways (Wnt, negative-regulation-of-Wnt, FGFR, ERK1/2, Notch, and EGFR.)

### Ethical approval

All methods were performed in accordance with the relevant guidelines and regulations. Ethical approval to access de-identified data from the UK Biobank and ABCD was obtained from the Human Biology Research Ethics Committee, University of Cambridge (Cambridge, UK) (HBREC.2020.07). Informed consent was provided by all participants.

## Data availability

GWAS summary statistics will be made available at the time of publication.

## Code Availability

All scripts used for analysis in this study were written in shell and R, and are available at https://github.com/yg330/GWAS <u>M</u>RI. The imaging pipeline used to generate are provided here: https://github.com/ucam-department-of-psychiatry/UKB. https://github.com/yg330/GWAS_MRI.

## Author contributions

YG and VW designed the study,conducted the primary analyses, and wrote the first draft of the manuscript. Additional analyses were conducted by AE and YH. RAIB and RRG provided quality controlled neuroimaging data for both cohorts. HW and HL provided data for MPRA analyses. CP, KK, SJ, ETB advised on various aspects of the study. All authors read the manuscript and provided critical feedback.

## Acknowledgements

VW receives funding from the European Unions Horizon 2022 R2D2-Mental Health project, SFARI, the MRC (MR/Z50354X/1), and the Wellcome Trust (309245/Z/24/Z and 214322\Z\18\Z). VW and EB are funded by the Medical Research Council ImmunoMIND hub (MR/Z50354X/1), part of the UKRI Mental Health Platform. RRG is funded by the Plan de Generación de Conocimiento from the Spanish Agencia Estatal de Investigación (PID2021-122853OA-I00 and PID2025-167805OB-I00), and ERANET Neuron JTC 2023 (ERP-2023-23684211). All research at the Department of Psychiatry in the University of Cambridge is supported by the NIHR Cambridge Biomedical Research Centre (BRC-1215-20014) and NIHR Applied Research Centre. The views expressed are those of the author(s) and not necessarily those of the NIHR or theDepartment of Health and Social Care. This work was supported by the John Lambton Trust fund and by the K. Lisa Yang Centre for Autism Research at Cambridge. Any views expressed are those of the author(s) and not necessarily those of the funders. The funders had no role in the design of the study, in the collection, analyses, or interpretation of data, in the writing of the manuscript, or in the decision to publish the results. For the purpose of Open Access, the authors have applied a CC BY public copyright licence to any Author Accepted Manuscript version arising from this submission.

