## Supplementary Figures for "Polygenic hierarchies of macroscale brain structural organisation"

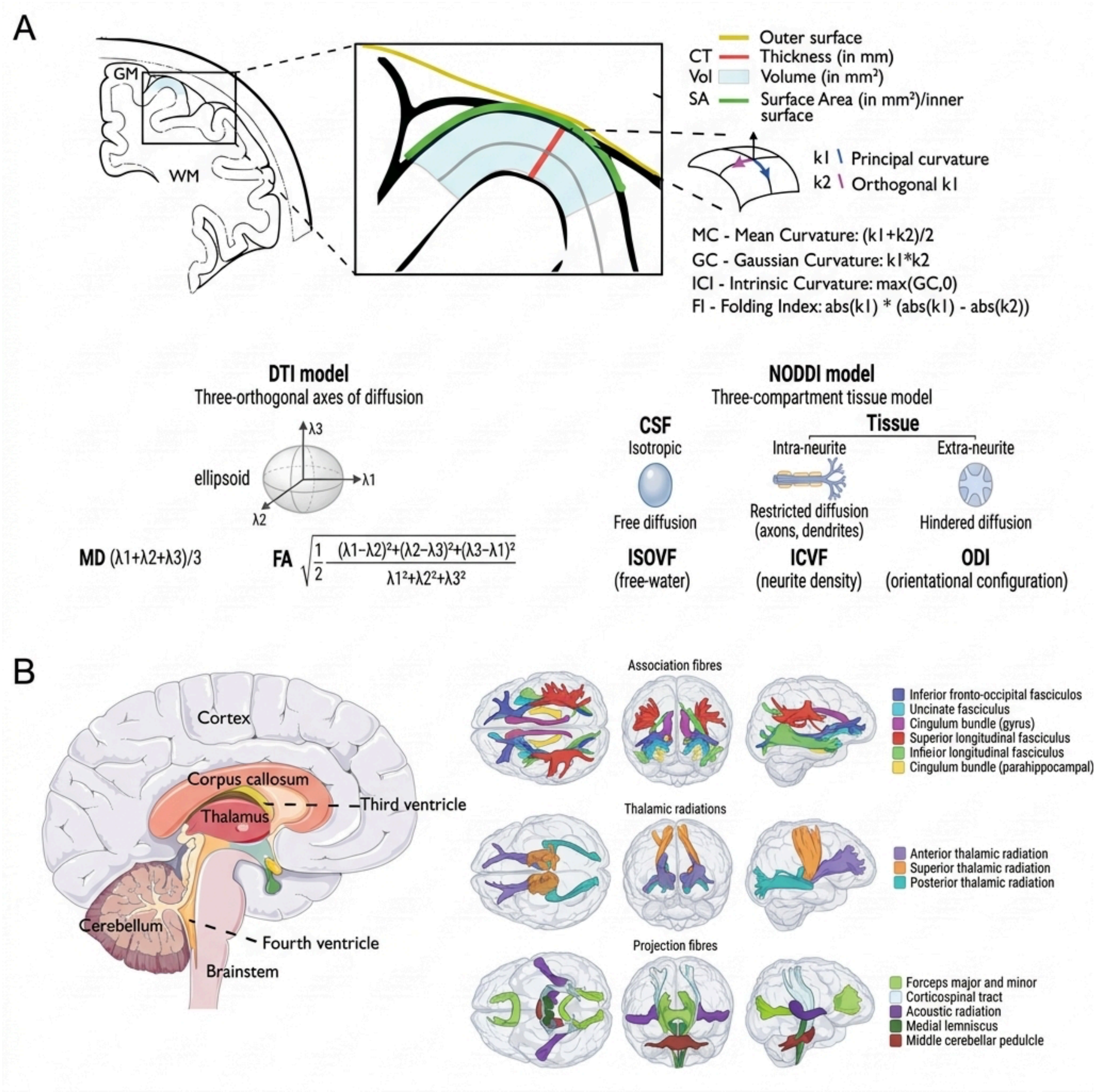


**Supplementary Figure 1: Overview of MRI-derived structural phenotypes and neuroanatomical structures analysed.** (**A**) Schematic showing the geometric basis of the structural measures used throughout: cortical thickness (CT), surface area (SA), and volume (Vol) from the pial/white matter surfaces; curvature measures derived from the principal curvatures k1 and k2 (mean curvature MC, Gaussian curvature GC, intrinsic curvature index ICI, folding index FI); and diffusion measures from the tensor (MD, FA) and NODDI (ISOVF, ICVF/NDI, ODI) models. (**B**) Representative schematic of the subcortical, periventricular, and white matter structures assessed, including major tract groups (association fibres, thalamic radiations, projection fibres) reconstructed via probabilistic tractography. This panel is illustrative rather than exhaustive and does not depict every subcortical or ventricular region included in the GWAS (Methods). Tract diagram in panel B adapted from Cox et al. (2016), https://doi.org/10.1038/ncomms13629, distributed under CC BY 4.0.

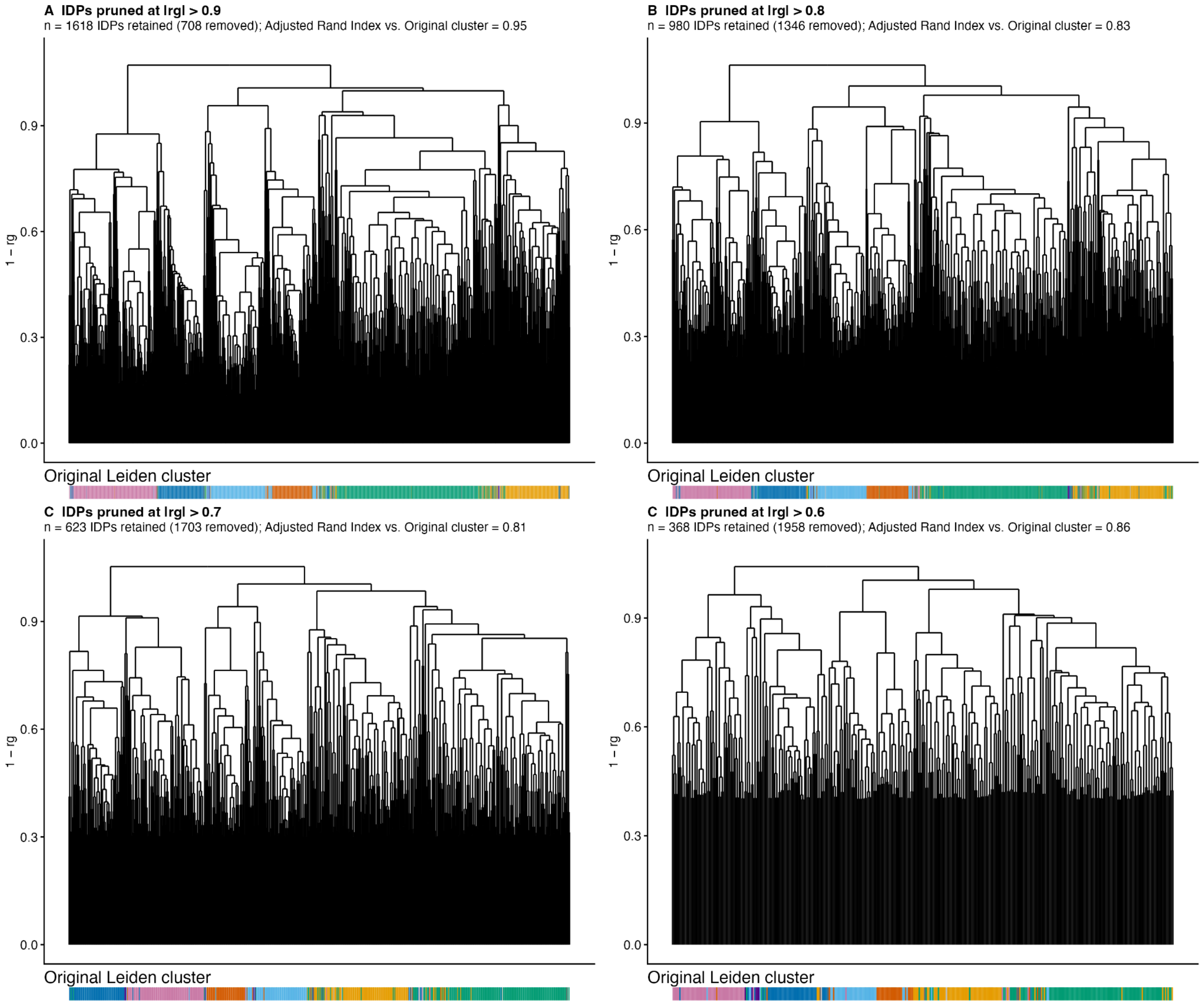


**Supplementary Figure 2: Consistency of hierarchical clustering after removing highly genetically correlated IDPs.** Average-linkage based hierarchical clustering dendrogram of all IDPs by genetic correlation distance (1 − rg), with annotation bar indicating Leiden community membership. This has been done separately after pruning highly genetically correlated IDPs and retaining 1 from each pair at a genetic correlation threshold of 0.9 (**A**); 0.8 (**B**), 0.7 ( **C**) and 0.6 (**D**). Similarity with the original hierarchical clustering dendrogram has been calculated using Adjusted Rand Index.

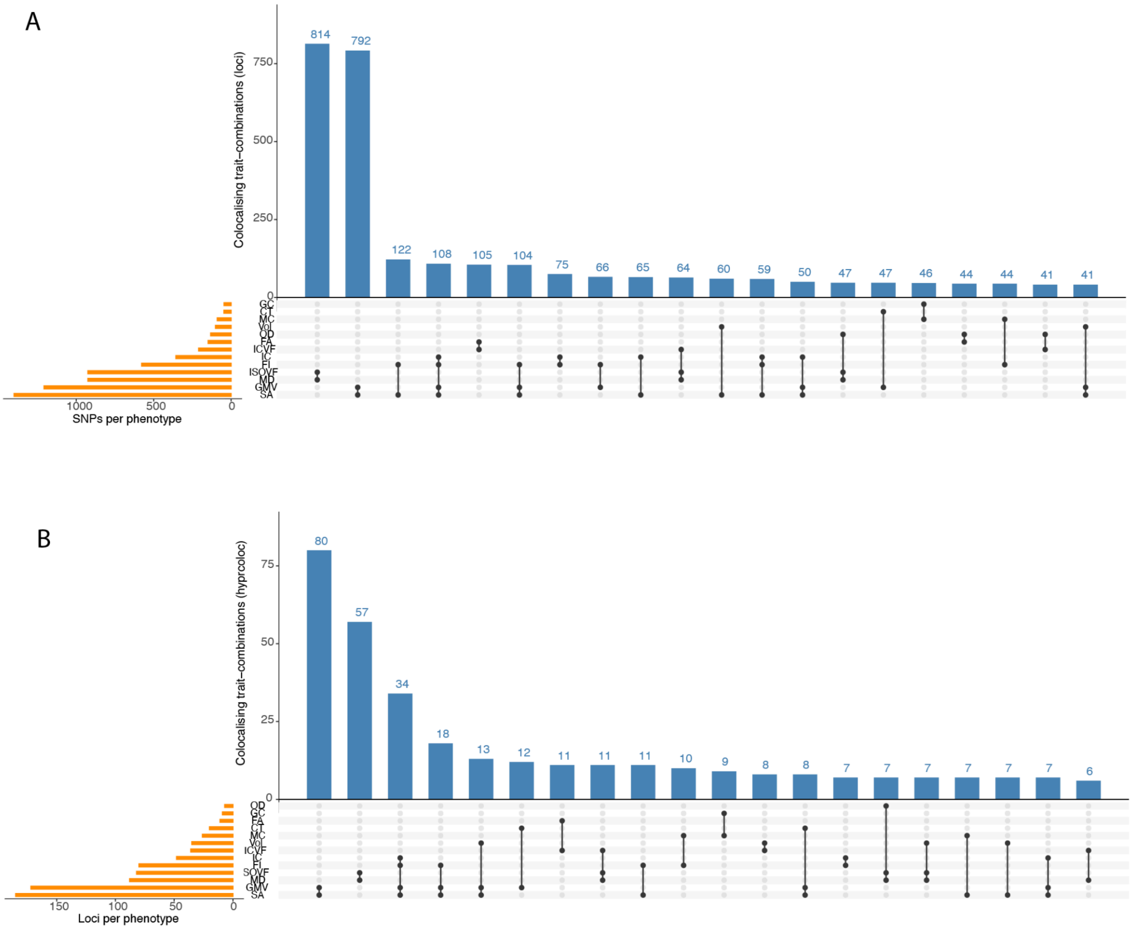


**Supplementary Figure 3. Overlap in shared loci between measures.** Upset plot showing overlap in experiment-wide significant loci after LD-based clumping (**A**) and colocalisation analysis using hyprcoloc (**B**).

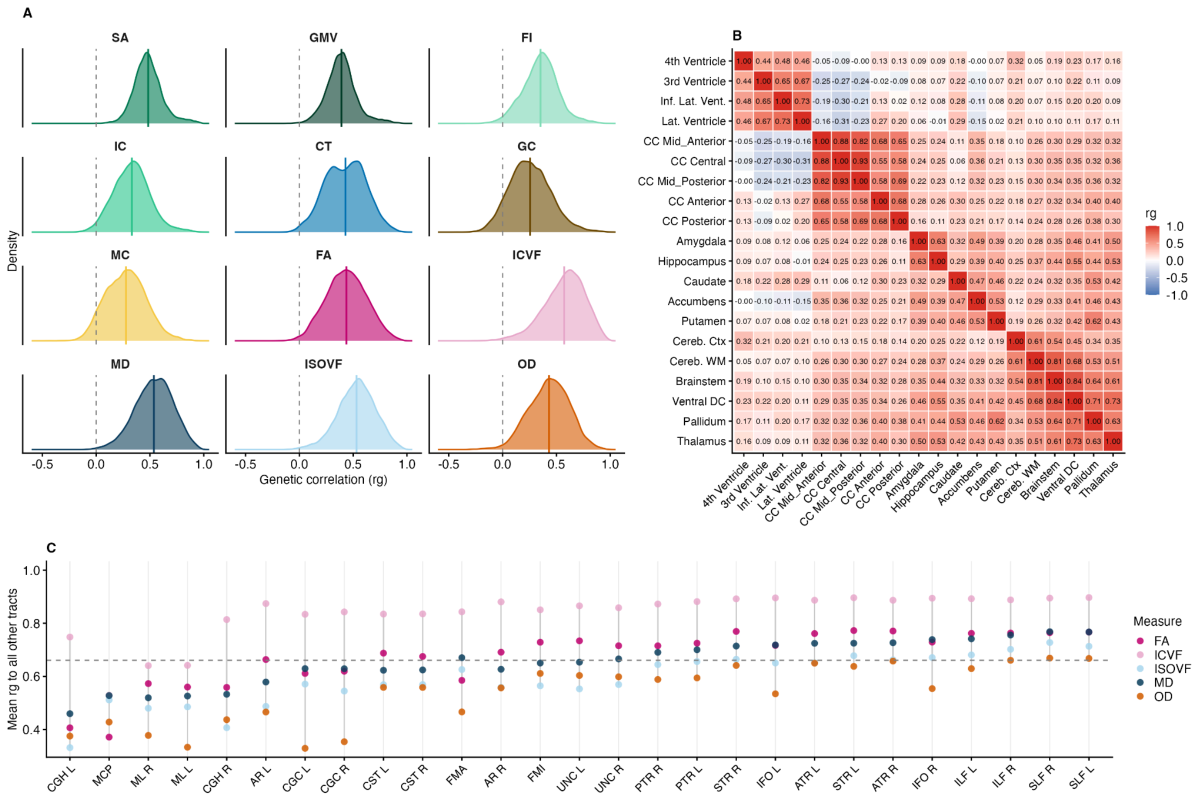


**Supplementary Figure 4. Fine-scale genetic architecture of structural MRI phenotypes. A.** Kernel density plots of pairwise within-measure genetic correlations across all 180 cortical regions, shown separately for each of the 12 structural MRI measures. Vertical lines indicate the mean genetic correlation per measure. **B.** Genetic correlation heatmap among subcortical, ventricular, and corpus callosal volumes. **C.** Mean genetic correlation of each white matter tract to all other tracts, calculated and visualised separately for each of the five microstructural measures (FA, ICVF, ISOVF, MD, OD). The dashed line indicates the median genetic correlation across all tract-measure combinations.

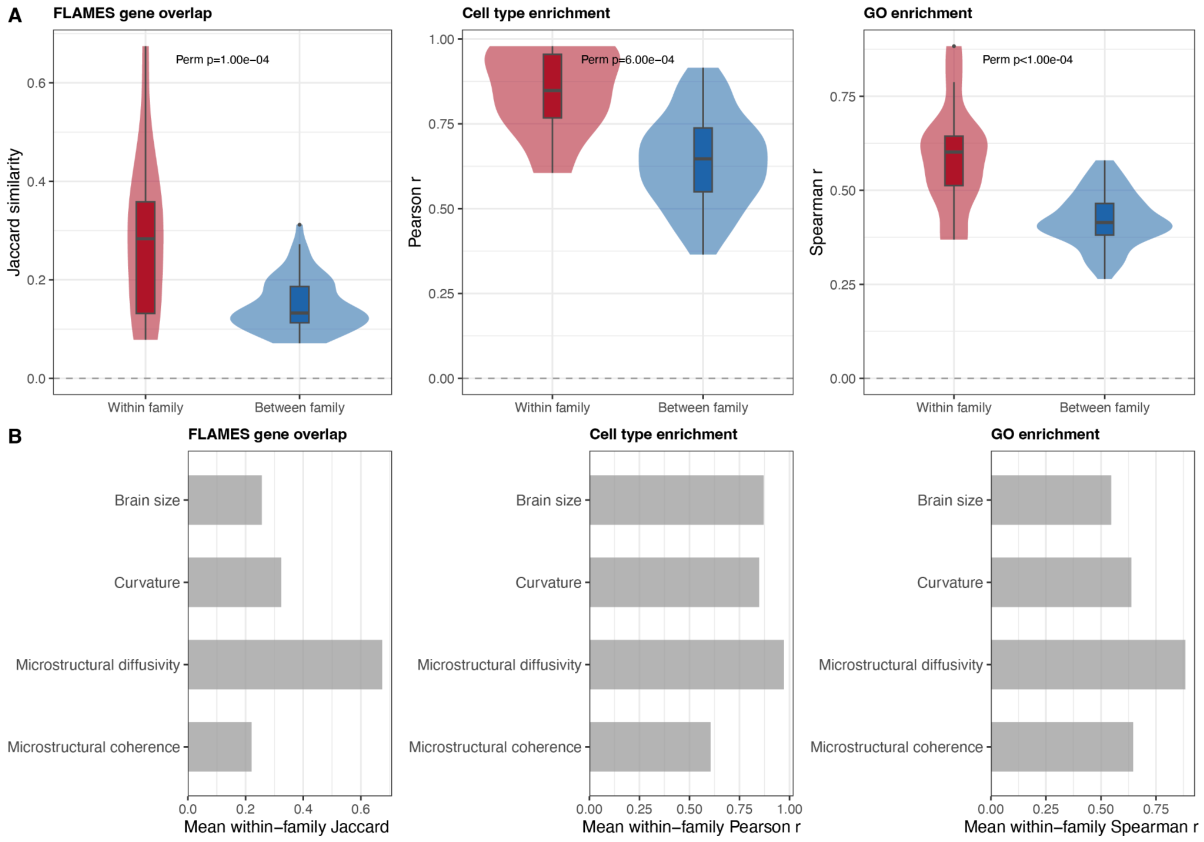


**Supplementary Figure 5. Within- and between-family similarity of FLAMES gene sets, cell type and Gene Ontology enrichment profiles. A.** Violin plots showing pairwise Jaccard index (FLAMES genes), Pearson correlation (cell type enrichment profiles), and Spearman correlation (Gene Ontology enrichment profiles) for within-family (red) and between-family (blue) phenotype pairs. Annotated with permutation p-value (n = 10,000). **B.** Mean within-family similarity per family for FLAMES, cell type, and GO analyses.

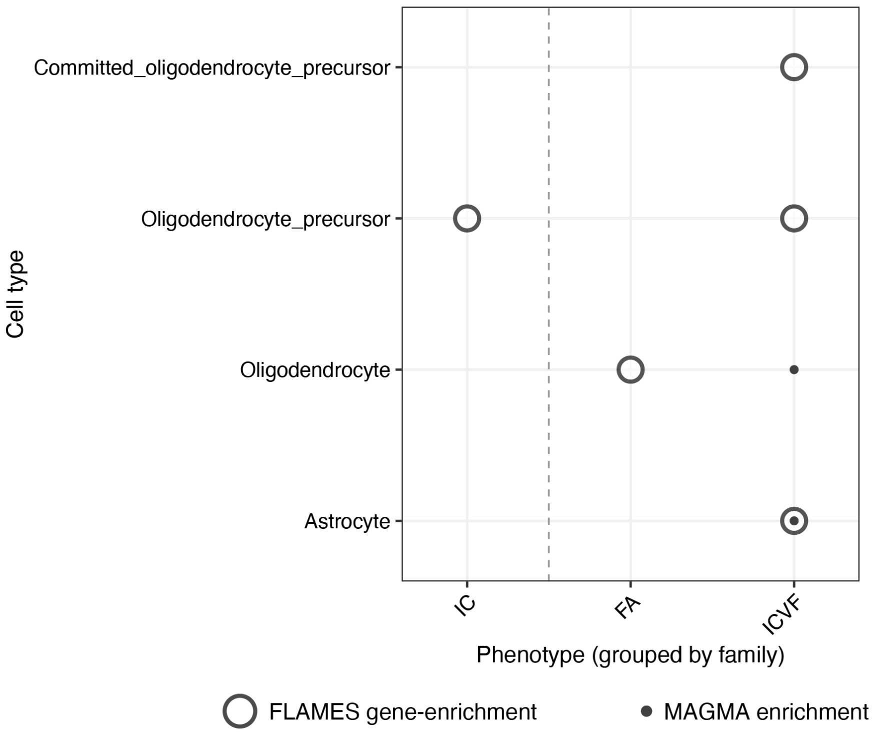


**Supplementary Figure 6: Significant cell type enrichment of FLAMES genes across Siletti 2023 adult cell types.** Outer circle indicates enrichment for FLAMES genes (logistic regression FDR < 0.05, $\beta > 0$). Filled dark grey inner circles indicate FDR-significant enrichment in the Wang 2025 MAGMA analysis of global GWAS signals ($\beta > 0$). Phenotypes are grouped by family (dashed vertical lines).

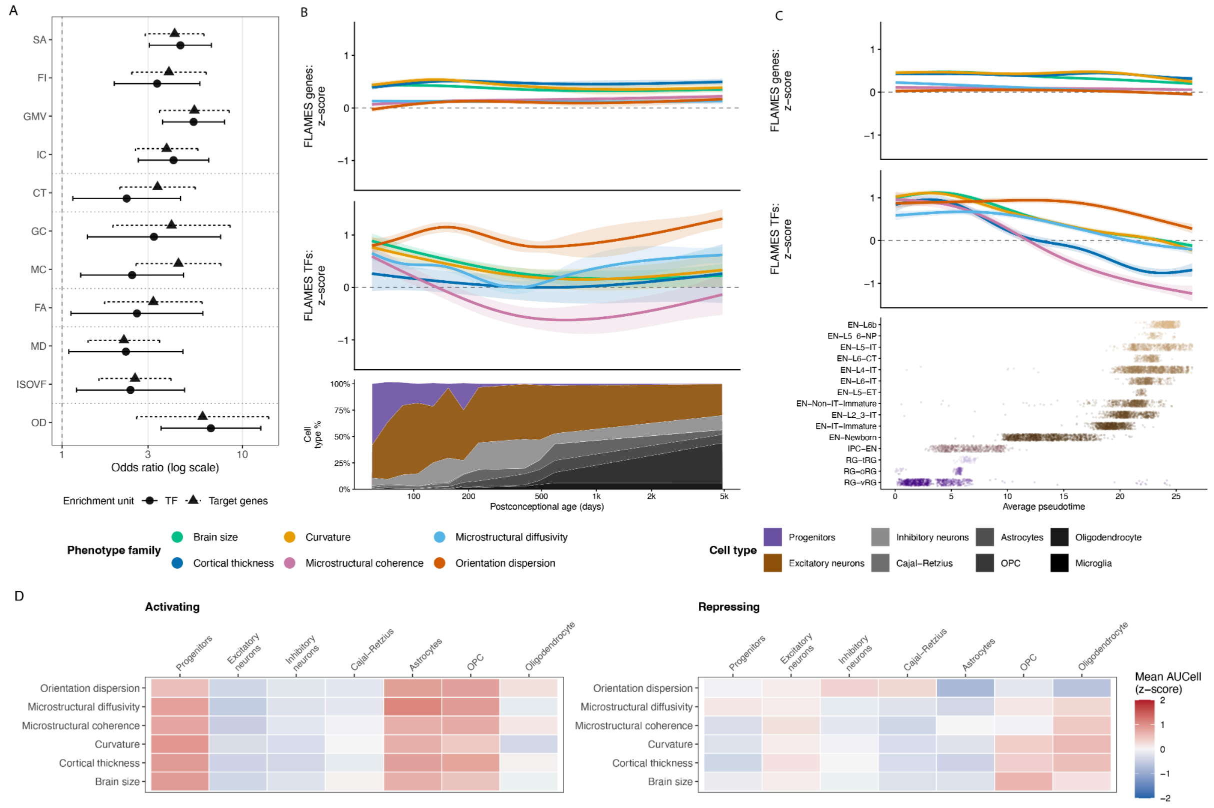


**Supplementary Figure 7. Gene regulatory mechanisms underlying brain structure. A.** Enrichment of eRegulon transcription factors (TFs, circles) and their target genes (triangles) among FLAMES-prioritised genes per phenotype. Points, odds ratio; error bars, 95% CI; only Benjamini-Hochberg FDR-significant associations shown. **B**. Relative expression (z-score against the whole transcriptome at each timepoint) of all significant FLAMES genes (top) and FLAMES-prioritised transcription factors (middle) across chronological postconceptional age, coloured by phenotype family; shaded bands, 95% CI. Bottom, proportion of major cell types across the same age axis. **C.** As in **b**, but along average pseudotime for the radial glia (RG) to excitatory neuron (EN) trajectory; the bottom panel instead shows individual cells per fine-grained RG/IPC/EN subtype, coloured by lineage. **D**. Mean AUCell activity (z-score) of activating (left) and repressing (right) eRegulons, averaged by phenotype family and broad cell type across all FDR-significant FLAMES-overlapping eRegulons.

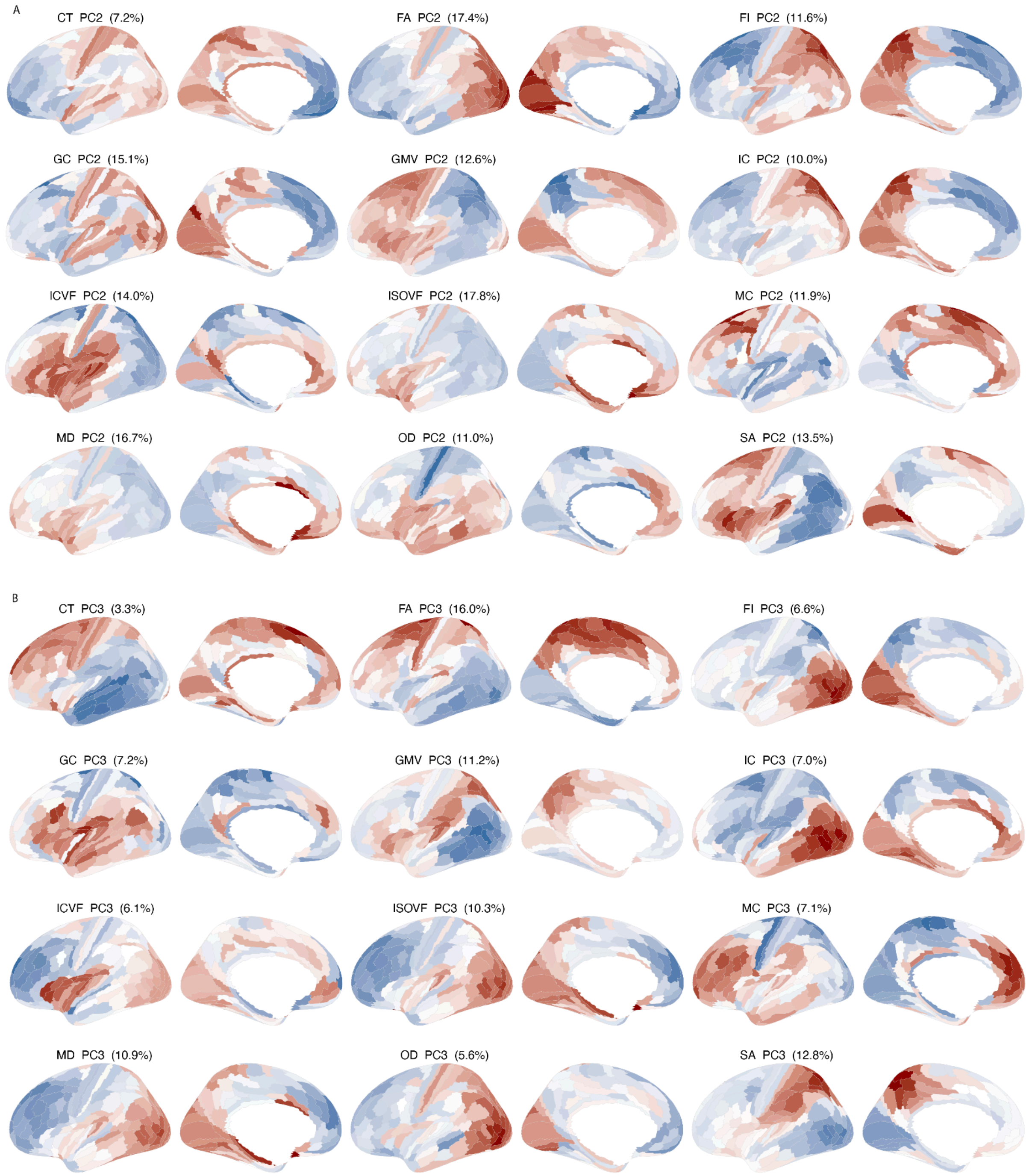


**Supplementary Figure 8. Spatial projection of the second two principal components on the Glasser parcellation by measure. A.** Spatial projection of gPC2**. B.** Spatial projection of gPC3

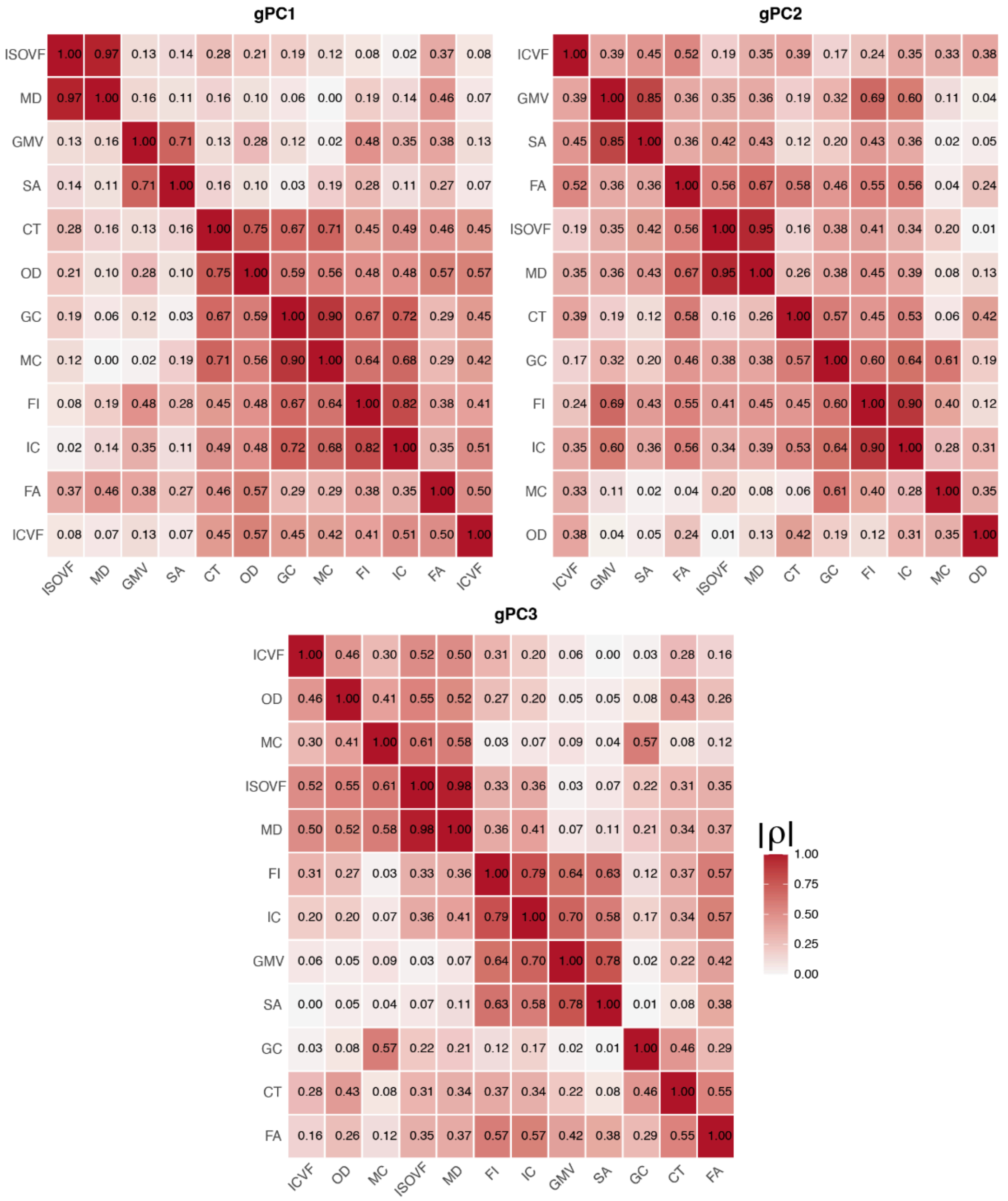


**Supplementary Figure 9. Cross-measure correspondence of gPC1–3.** For each of the top three genetic principal components (gPC1, gPC2, gPC3) derived independently for each of the 12 structural measures, heatmaps show the absolute Spearman correlation (|ρ|) between measures' parcel-wise component loadings, computed pairwise across all 12 measures. Component sign is arbitrary (Methods), so |ρ| is shown rather than signed ρ. Within each panel, measures are ordered by hierarchical clustering (average linkage, distance = 1 − |ρ|) of that component's own similarity structure; the resulting order can therefore differ between gPC1, gPC2, and gPC3.

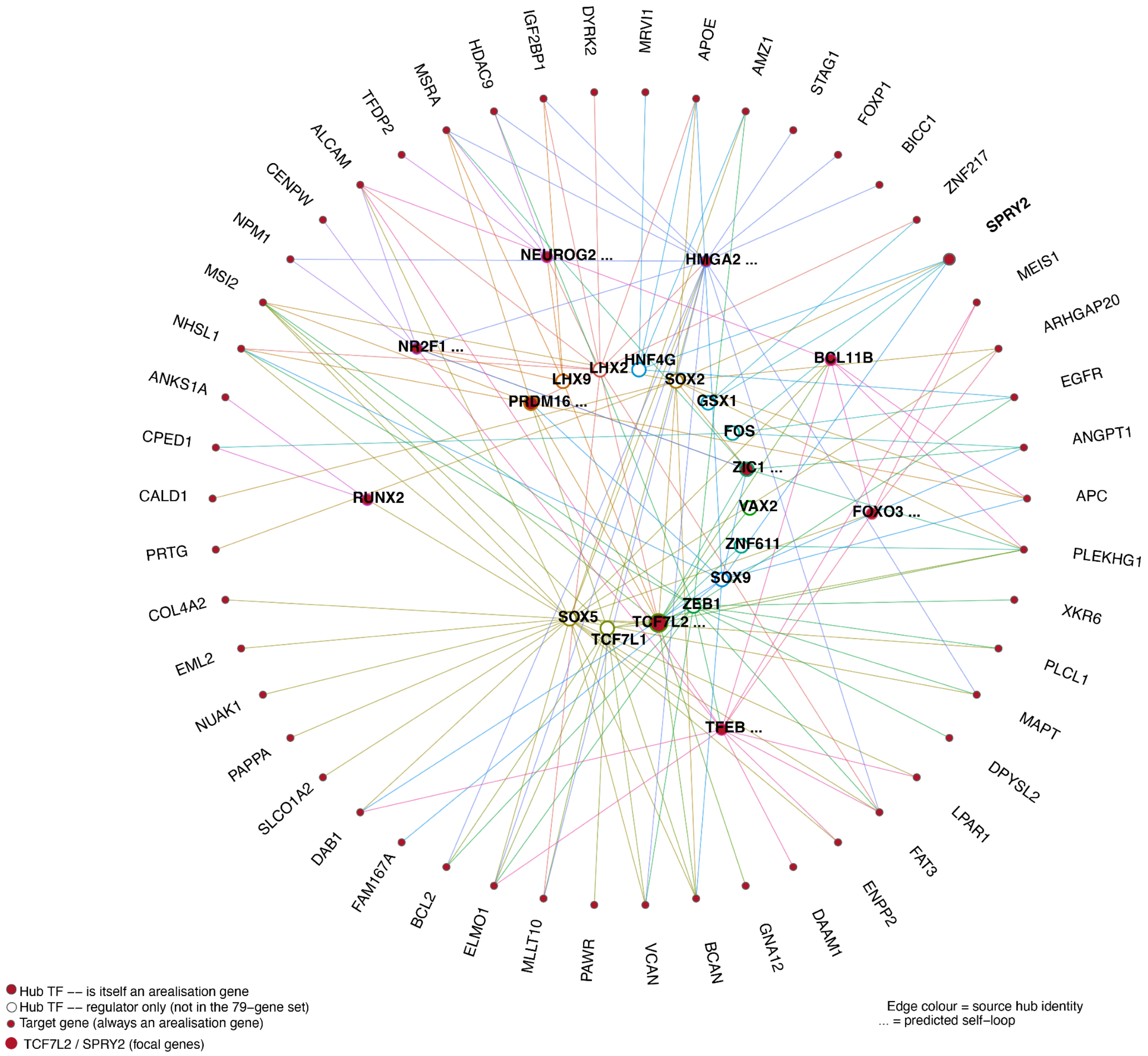


**Supplementary Figure 10. Gene regulatory network (eGRN) underlying TCF7L2 and SPRY2.** Radial network showing predicted transcriptional regulators of TCF7L2 (a Wnt-pathway effector) and SPRY2 (an FGF-pathway antagonist), both identified as arealization genes in this study. Inner ring are transcription factors whose predicted eRegulon target set is significantly enriched for TCF7L2 and/or SPRY2 (FDR<0.05,). Middle ring are TFs found among the inner-ring hubs' target genes but are arealization genes themselves with their own predicted targets. Target genes (outer ring) are each hub's predicted eRegulon targets, restricted to the 79 arealization genes identified in this study. Edge colour indicates the source (regulating) hub TF. Node fill: maroon indicates the gene is itself one of the 79 arealization genes; white indicates a TF identified purely as a regulator, not independently detected as an arealization gene in the GWAS.
